# Immunodominant extracellular loops of *Treponema pallidum* FadL outer membrane proteins elicit antibodies with opsonic and growth-inhibitory activities

**DOI:** 10.1101/2024.07.30.605823

**Authors:** Kristina N. Delgado, Melissa J. Caimano, Isabel C. Orbe, Crystal F. Vicente, Carson J. La Vake, André A. Grassmann, M. Anthony Moody, Justin D. Radolf, Kelly L. Hawley

## Abstract

The global resurgence of syphilis has created a potent stimulus for vaccine development. To identify potentially protective antibodies (Abs) against *Treponema pallidum* (*TPA*), we used *Pyrococcus furiosus* thioredoxin (*Pf*Trx) to display extracellular loops (ECLs) from three *TPA* outer membrane protein families (outer membrane factors for efflux pumps, eight-stranded β-barrels, and FadLs) to assess their reactivity with immune rabbit serum (IRS). Five ECLs from the FadL orthologs TP0856, TP0858 and TP0865 were immunodominant. Rabbits and mice immunized with these five *Pf*Trx constructs produced ECL-specific Abs that promoted opsonophagocytosis of *TPA* by rabbit peritoneal and murine bone marrow-derived macrophages at levels comparable to IRS and mouse syphilitic serum. ECL-specific rabbit and mouse Abs also impaired viability, motility, and cellular attachment of spirochetes during *in vitro* cultivation. The results support the use of ECL-based vaccines and suggest that ECL-specific Abs promote spirochete clearance via Fc receptor-independent as well as Fc receptor-dependent mechanisms.

**Author Summary:** The resurgence of syphilis emphasizes the critical need for vaccine development against *Treponema pallidum* (*TPA*). Research utilizing immune rabbit serum (IRS) suggests that an effective syphilis vaccine should induce “functional” antibodies (Abs) capable of enhancing the opsonophagocytosis of treponemes by activated macrophages. Structural models of *TPA* outer membrane proteins (OMPs), specifically the extracellular loops (ECLs), guided the identification of potential vaccine candidates. Antigenic analysis with IRS of individual ECLs from three *TPA* OMP families scaffolded onto *Pyrococcus furiosus* thioredoxin (*Pf*Trx) revealed five FadL antigenic ECLs. Immunization with immunodominant ECL antigens elicited robust ECL-specific Abs, demonstrating functional opsonic activity in the opsonophagocytosis assays. Furthermore, these Abs effectively inhibited the growth inhibition of *in vitro*-cultivated *TPA* in both rabbit and mouse models. Our findings underscore the value of antigenic analysis in identifying promising *TPA* OMP ECL vaccine targets and highlight the multifaceted protective capacity of ECL Abs against *TPA*. This approach also extends to identifying potential OMP vaccinogens in other bacterial pathogens, offering valuable insights for broader vaccine development strategies.

## Introduction

Syphilis is a multistage, sexually transmitted infection caused by the highly invasive and immunoevasive spirochete *Treponema pallidum* subsp. *pallidum* (*TPA*)[1, 2]. Since the start of the current millennium, the disease has undergone a dramatic resurgence in the United States and worldwide even though its causative agent remains exquisitely susceptible to penicillin after more than seven decades of use[1–3]. These alarming trends underscore the urgent need for new control strategies, including vaccines[4, 5]. The rabbit has long been considered the animal model of choice for investigating protective immunity against syphilitic infection[6–8]. Rabbits develop long-lasting immunity to reinfection[6–9], and it is generally believed that deconvolution of protective responses in the rabbit will inform vaccine development for humans. Evidence from the rabbit model[10], supported by subsequent *ex vivo* studies with human syphilitic sera[11–13], has brought to light the importance of macrophage-mediated opsonophagocytosis as a primary mechanism for clearance of *TPA*. Accordingly, it is generally believed that opsonic antibodies (Abs) for *TPA* can be considered a surrogate for protection[10, 14]. Whether opsonophagocytosis is the sole mechanism for Ab-mediated clearance of *TPA* in humans or animals, however, remains to be determined. Historically, mouse models have not found widespread acceptance in the syphilis field[15, 16]. Nevertheless, *TPA*-infected mice clear the infection and produce Abs that promote uptake and degradation of spirochetes by bone marrow-derived macrophages (BMDMs)[17–19]. These results suggest that the mouse model has potential utility to expedite selection and evaluation of syphilis vaccine candidates.

Extensive investigation of the molecular architecture of the *TPA* outer membrane (OM) has identified the spirochete’s repertoire of OM proteins (OMPs) as the principal candidate antigens for syphilis vaccine design[20–25]. The *TPA* OMPeome consists of two proteins, BamA (TP0326) and LptD (TP0515), involved in OM biogenesis and four paralogous families involved in importation of nutrients or extrusion of noxious substances across the OM: OM factors (OMFs) for efflux pumps, eight-stranded β-barrels (8SβBs), long-chain fatty acid transporters (FadLs), and *Treponema pallidum* repeat proteins (Tprs)[23, 24]. As in other diderm bacteria[26], the OM-embedded portions of *TPA* OMPs adopt a β-barrel conformation in which extracellular loops (ECLs) bridge neighboring β-strands[23, 24, 27]. So-called ‘functional’ Abs must target ECLs, the only Ab-accessible regions of OMPs, to promote clearance of spirochetes.

To study the antigenic properties of individual ECLs in a conformationally native-like state, they must be tethered, typically done using protein scaffolds[28–30]. We recently described use of *Pyrococcus furiosus* thioredoxin (*Pf*Trx)[31] as a scaffold for assessing the reactivity of *TPA* OMP ECLs with syphilitic sera and generating ECL-specific, opsonic Abs[18, 27]. Herein, we used immune rabbit sera (IRS) to assess the immunogenicity of scaffolded ECLs from three newly discovered OMP paralogous families: OMFs, 8SβBs, and FadLs[23, 24, 27]. With this strategy, we identified five immunodominant ECLs from three members of the FadL family and used *Pf*Trx-scaffolded ECLs to generate opsonic Abs in rabbits and mice. By exploiting the recent breakthrough in long-term *in vitro* cultivation of *TPA*[32], we discovered that rabbit and mouse opsonic Abs against immunodominant FadL ECLs affected, to varying extents, spirochete viability, motility, and attachment to rabbit epithelial cells. Notably, removal of immune pressure by passage of organisms into Ab-free medium substantially rescued spirochete growth and motility. Collectively, our findings support a strategy for syphilis vaccine development based upon targeting of ECLs, and they provide novel insights into the mechanisms whereby Abs against *TPA* surface epitopes promote spirochete clearance.

## Results

### Prediction of ECL boundaries using structural models generated by trRosetta and AlphaFold3

Previously, we used trRosetta[33] to generate three-dimensional (3D) structural models for three recently discovered *TPA* OMP paralogous families: OMFs, 8SβBs, and FadLs (**Fig 1** and **S1 Fig**)[23, 24, 27]. These 3D models enabled us to identify the putative ECL boundaries (**S1 Table**) needed to create *Pf*Trx-scaffolded ECLs for the antigenicity analyses described below. The subsequent emergence of AlphaFold3[34] as the leading program for 3D protein modeling prompted us to reanalyze the predicted ECL boundaries. High-confidence models from AlphaFold3 and trRosetta for the 8SβBs and FadL families demonstrated strong agreement of ECL boundaries (**S1A and S1C Fig**). FadL proteins contain an N-terminal extension (‘hatch’) that occludes the lumen of the β-barrel[35, 36]. A distinctive feature of the *TPA* FadL family is an N-terminal hatch predicted to extend through the β-barrel to the extracellular space[24]; notably AlphaFold3 and trRosetta predictions for the hatches were consistent for all five FadLs (**S1C Fig**). Three of the five FadL orthologs (TP0548, TP0859, and TP0865) feature α-helical C-terminal extensions, presumed to be periplasmic, also predicted by both AlphaFold3 and trRosetta. Predictions by trRosetta and AlphaFold3 for the *TPA* OMFs, however, diverged substantially (**S1A Fig** and **S2 Fig**). Structurally characterized OMFs are homotrimers in which the monomers contain four β-strands, two ECLs, and six extended α-helices (see examples *E. coli* TolC and *Neisseria gonorrhoeae* MtrE in **S2 Fig**)[37, 38]. trRosetta predicts canonical monomeric structures for all four *TPA* OMFs (TP0966, TP0967, TP0968, and TP0969). In contrast, AlphaFold3 models each monomer with eight β-strands, four small ECLs, and six α-helices. Based on the many solved structures available, we concluded that the trRosetta prediction of ECL boundaries are more likely to be correct and, therefore, used the four β-stranded monomer to complete the trimeric models using WinCoot[24] (**Fig 1A**).

**Fig 1.**
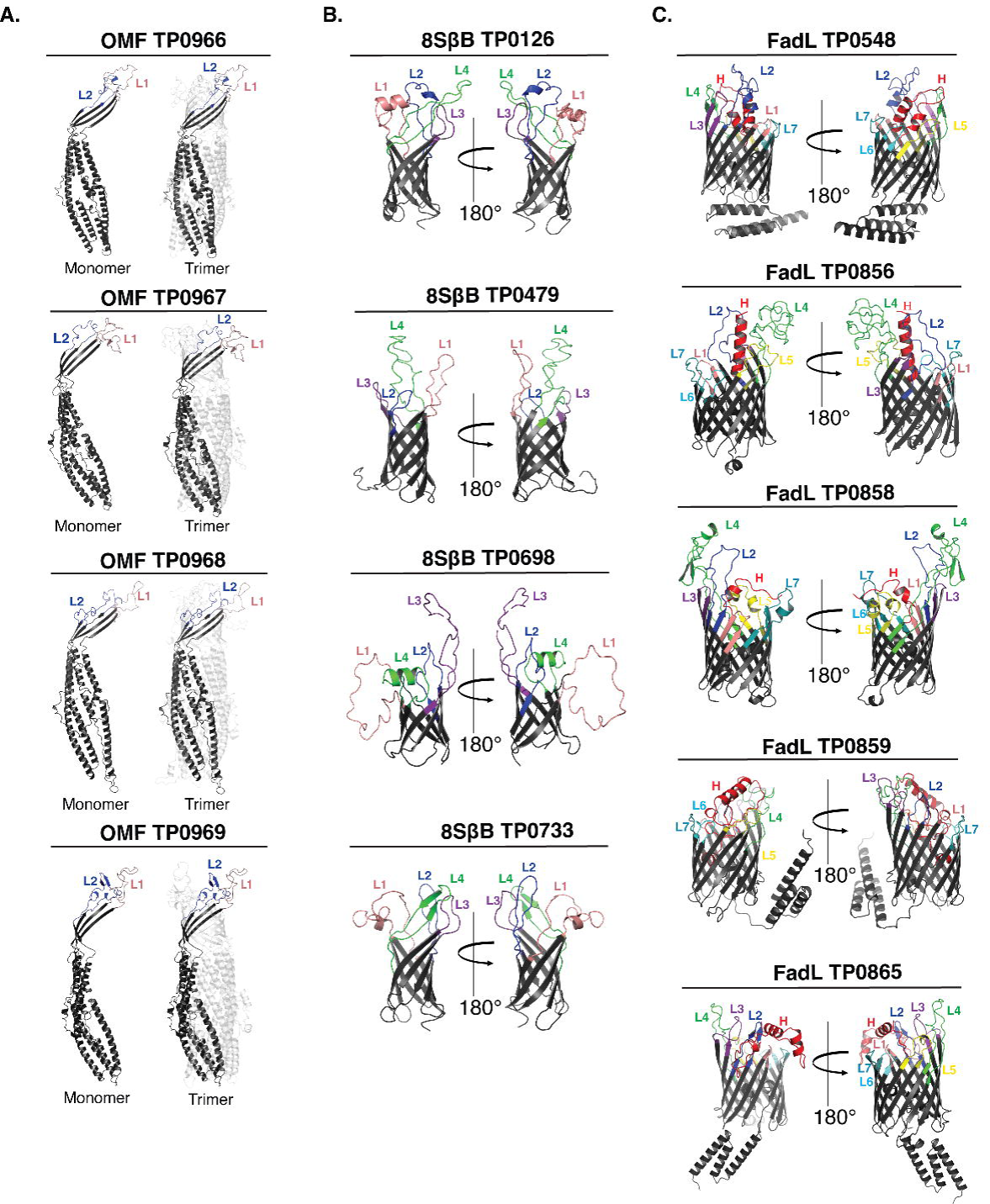
Prediction of ECL boundaries. trRosetta 3D models for outer membrane factors (OMFs), eight-stranded β-barrels (8SβBs), and FadLs (Panels A-C, respectively), depict ECL boundaries (ECL1-Salmon, ECL2-Blue, ECL3-Purple, ECL4-Green, ECL5-Yellow, ECL6-Cyan, ECL7-Dark Teal, and Hatch-Red) used to clone ECLs onto the *Pf*Trx scaffold (see **S1 Table**).

### Predicted linear and discontinuous B cell epitopes reside predominantly in ECLs

As a starting point for our analysis of ECL reactivity with IRS, we used ElliPro[39] and DiscoTope 2.0[40] to predict BCEs across the three paralogous families. As shown in Figure 2 and Supporting Figure 3, the predictions for linear and discontinuous BCEs mapped predominantly to ECLs. Notably, only ElliPro predicted discontinuous epitopes for ECL2 of TP0967 and TP0968 (**Fig 2A** and **S3A Fig**). For the 8SβBs, ECL2 and ECL3 of TP0479 lacked predicted linear and conformational BCEs (**Fig 2B** and **S3B Fig**). The FadL family presented a more complex picture. While most ECLs exhibited both linear and discontinuous epitopes, some ECLs (*e*.*g*., ECL1 of TP0548 and ECL7 of TP0865) were predicted to possess only linear epitopes (**Fig 3** and **S3C Fig**).

**Fig 2.**
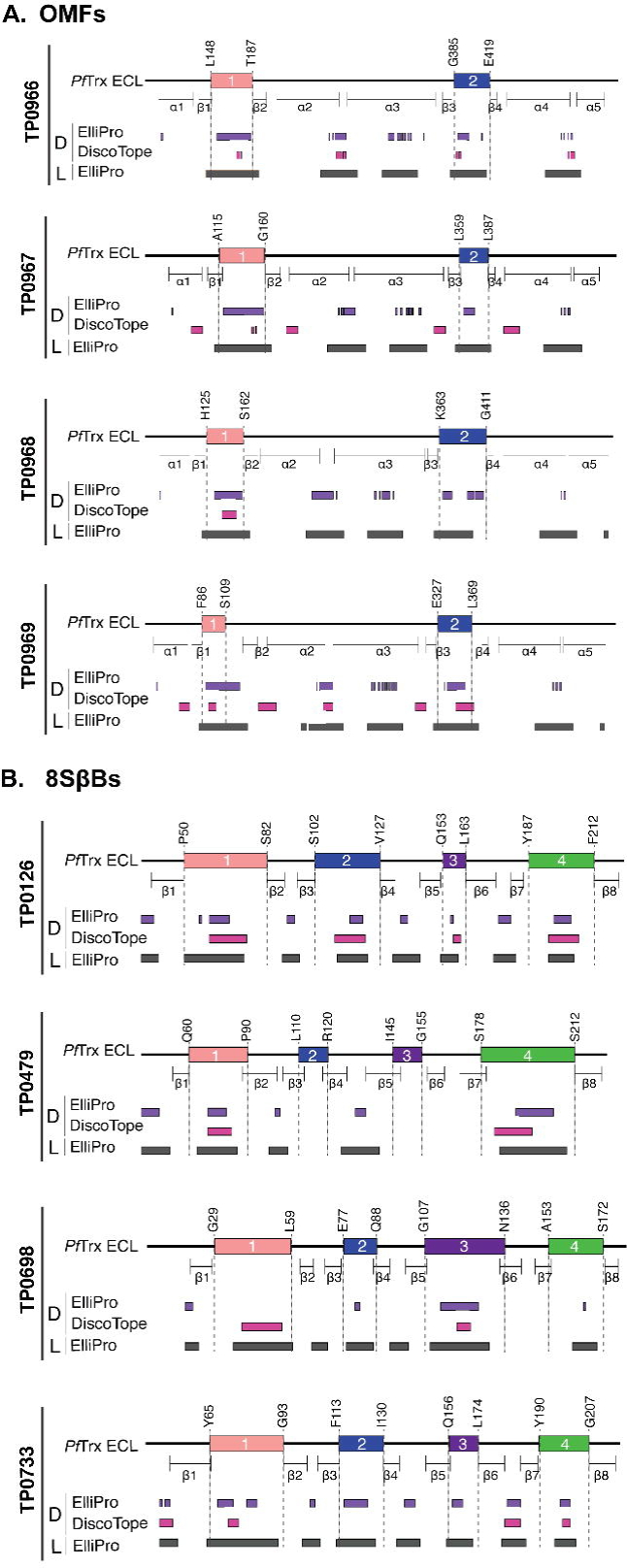
B cell epitope (BCE) predictions for OMFs and 8SβBs. One-dimensional (1D) models depicting the positions of linear (L) and discontinuous (D) BCEs predicted by ElliPro[39] (threshold ≥0.8) and DiscoTope[40] (threshold -3.7) for (**a**) OMFs and (**b**) 8SβBs. 1D models display *Pf*Trx-scaffolded ECLs using the color scheme described in the caption for Figure 1.

**Fig 3.**
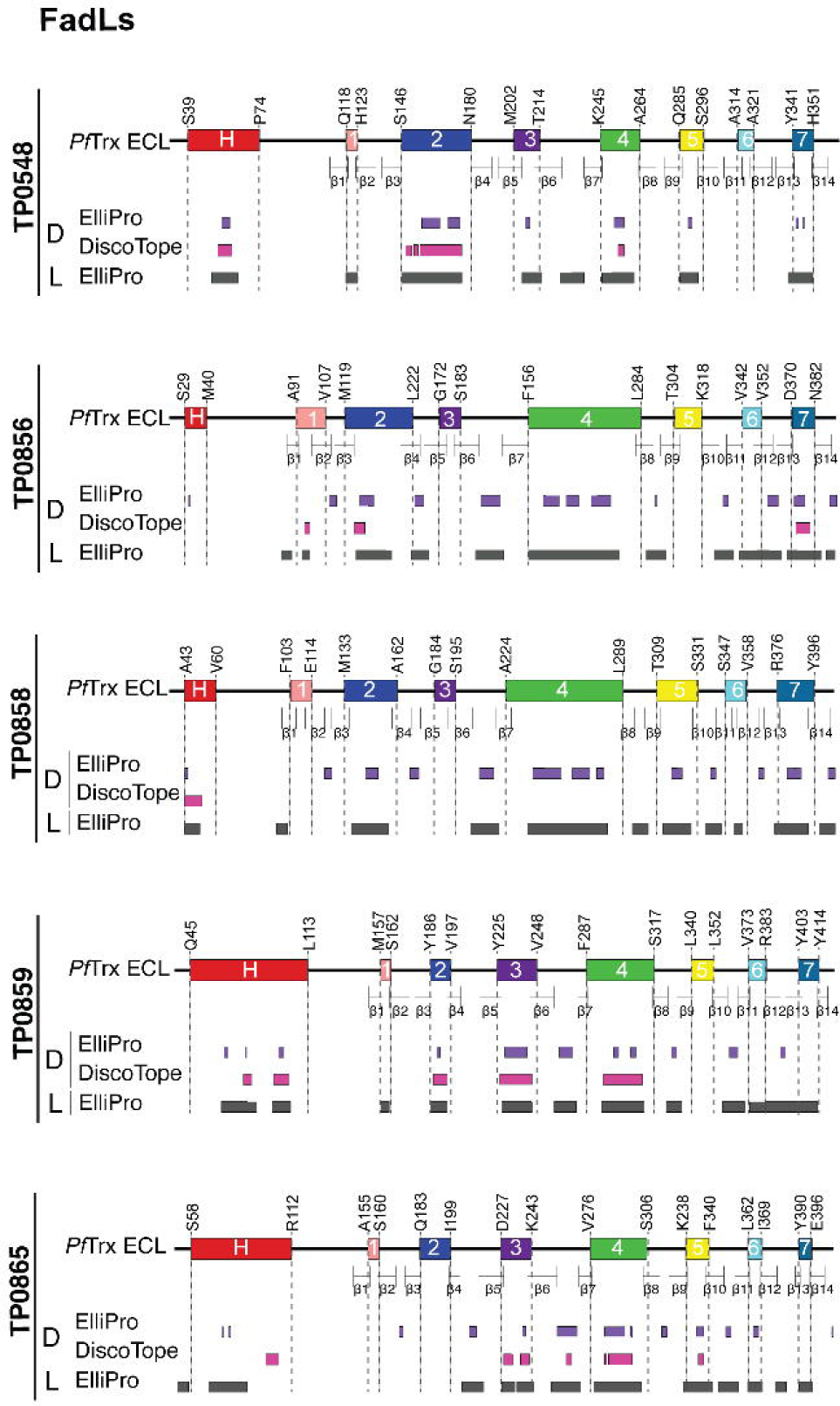
B cell epitope (BCE) predictions for FadLs. One-dimensional (1D) models depicting the positions of linear (L) and discontinuous (D) BCEs predicted by ElliPro[39] (threshold ≥0.8) and DiscoTope[40] (threshold -3.7) for the FadLs. 1D models display *Pf*Trx-scaffolded ECLs using the color scheme described in the caption for Figure 1.

### Antigenic analysis of scaffolded ECLs with *TPA* Nichols IRS reveals immunodominant FadL ECLs

Before proceeding to the examination of ECLs, we first assessed the reactivity of five Nichols IRS by immunoblotting against whole cell lysates from the same strain (**S4A Fig**). While each serum reacted strongly with known immunogenic lipoproteins (*e.g.,* Tpp47, Tpp17, and Tpp15)[41, 42], we noted differences in their recognition of other *TPA* proteins as would be expected for outbred animals. We then examined the reactivity of scaffolded OMF, 8SβB, and FadL ECLs with the immune sera by immunoblot and ELISA. Not surprisingly, the five immune rabbits exhibited considerable heterogeneity in Ab responses to ECLs of all three OMP families. Despite strong BCE predictions, the OMF and 8SβB ECLs showed poor reactivity overall (**Fig 4**). We also noted discordances between immunoblot and ELISA results for several OMF ECLs. For example, the strong ELISA reactivity of IRS 112 and 718 with both ECLs of TP0966 contrasted with their faint reactivity by immunoblot. Conversely, IRS 112 reacted strongly by immunoblot with ECL1 of TP0968 and ECL2 of TP0969 but showed no reactivity by ELISA (**Fig 4A**). Similar discordances were noted for the 8SβBs (**Fig 4B**). IRS 112 exhibited strong immunoblot reactivity for several 8SβBs ECLs that were non-reactive by ELISA, while IRS 113 reacted strongly by ELISA with ECL4 of TP0698 but showed no reactivity by immunoblot (**Fig 4B**). ECLs of the FadLs TP0856, TP0858, and TP0865 were the most immunoreactive overall (**Fig 5**). ECL2 and ECL4 of TP0856 and TP0858 along with ECL3 of TP0865 displayed strong reactivity by both immunoblot and ELISA. It is noteworthy that all five strongly reactive FadL ECLs were predicted to contain both linear and conformational BCEs. On the other hand, other FadL ECLs with strong BCE predictions were weakly antigenic.

**Fig 4.**
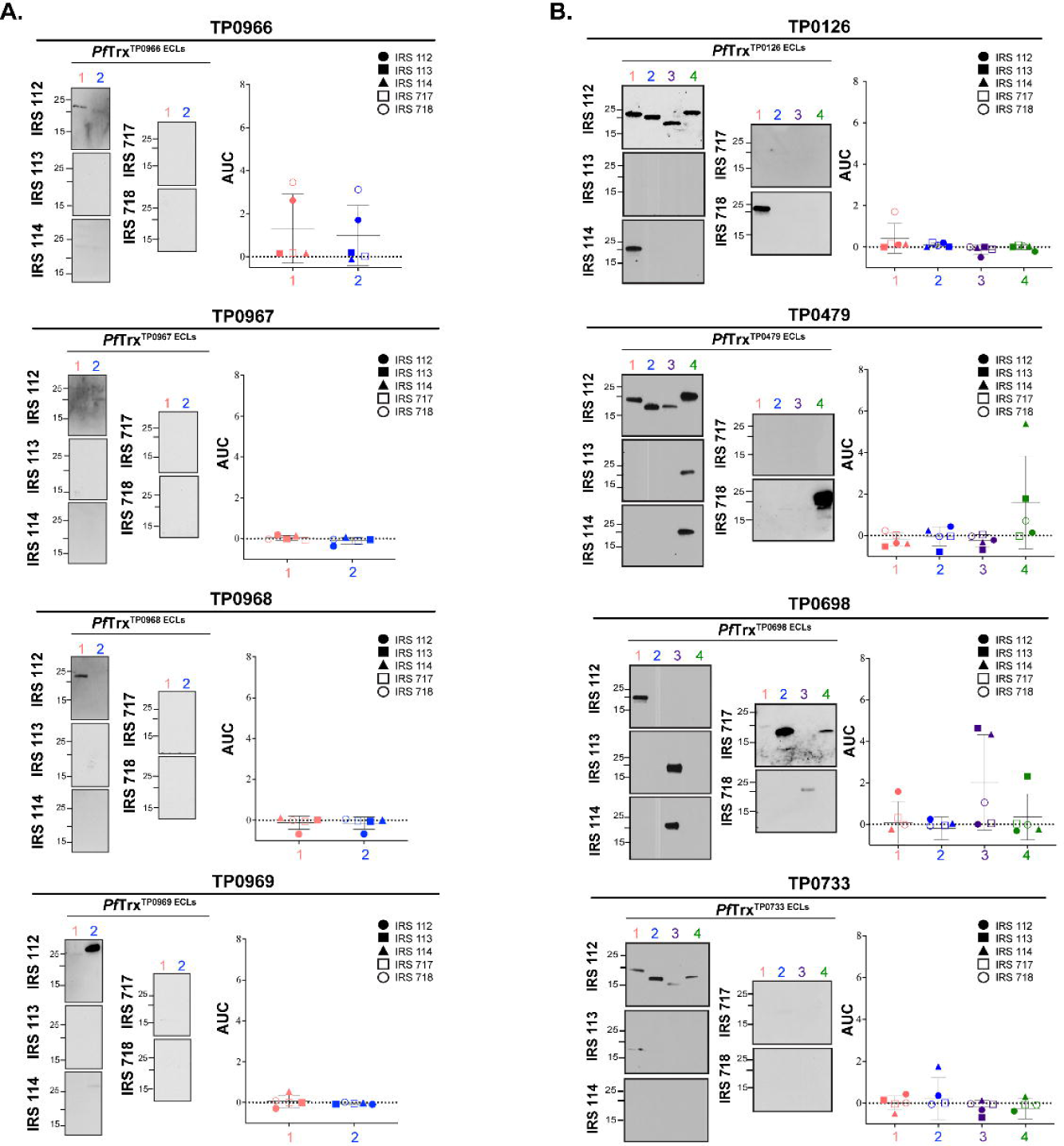
Reactivity of scaffolded ECLs with Nichols IRS reveals poorly immunogenic OMF and 8SβB ECLs. Reactivity by immunoblot (left) and ELISA (right) of scaffolded ECLs of (**A**) OMFs and (**B**) 8SβBs against sera from five Nichols immune rabbits. ELISA reactivity was measured as area under the curve (AUC) corrected for *Pf*Trx background (see Methods). *n*L=L3 wells per condition. Data are shown as meanL±LSD.

**Fig 5.**
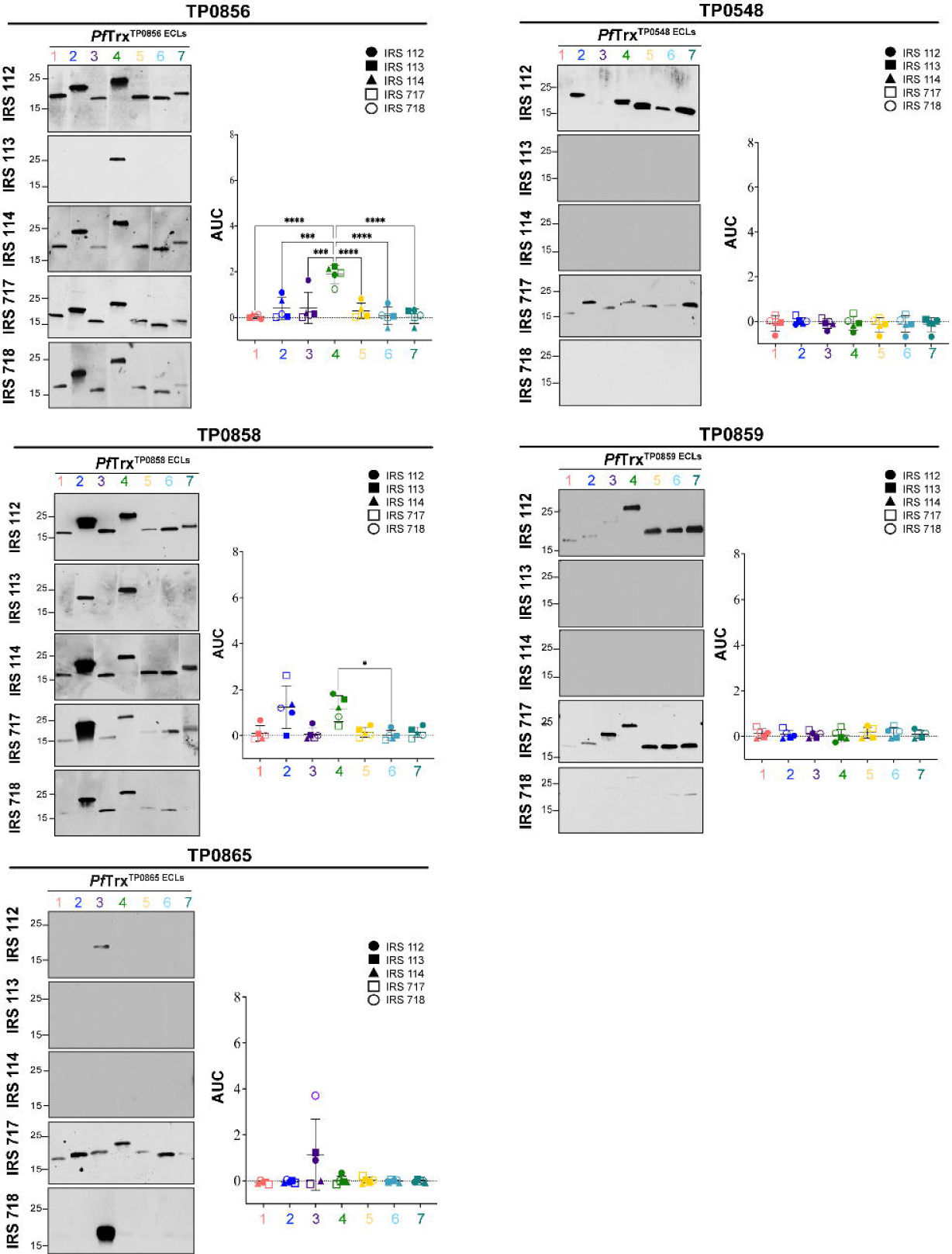
Reactivity of scaffolded ECLs with Nichols IRS reveals immunodominant FadL ECLs. Reactivity by immunoblot (left) and ELISA (right) of scaffolded FadL ECLs against sera from five Nichols immune rabbits. ELISA reactivity was measured as AUC corrected for *Pf*Trx background (see Methods). *n*L=L3 wells per condition. Data are shown as meanL±LSD. Significant differences (\**p*<0.05; \*\*\**p*<0.001; or \*\*\*\**p*<0.0001) between the means of the groups were determined by one-way ANOVA with Bonferroni’s correction for multiple comparisons.

Genomic sequencing has revealed that syphilis spirochetes cluster into two taxonomic groups represented by the Nichols and SS14 reference strains, with SS14-like strains predominating globally[43–45]. Given the epidemiologic importance of the SS14 clade, we next sought to determine whether SS14 immune rabbits also generate Abs against FadL ECLs. We first assessed the reactivity of three SS14 immune sera by immunoblotting against Nichols *TPA* lysates (**S4A Fig**). As with the Nichols IRS, SS14 immune sera reacted strongly with known immunogenic lipoproteins, although, once again, minor differences were noted in their recognition of other *TPA* proteins. Sequence alignment of the Nichols and SS14 FadL orthologs (**S5 Fig**) revealed that TP0856 and TP0859 are completely conserved. Compared to the Nichols ortholog, SS14 TP0858 harbors a single conservative amino acid substitution in ECL7 (S380N). SS14 TP0865 contains a non-conservative substitution in ECL2 (A193T) as well as an insertion of an asparagine residue at position 238 in ECL3. TP0548, on the other hand, displayed substantial variability in four ECLs (2, 4, 5 and 6). **Figure 6A** presents a summary of the variable residues within the Nichols and SS14 FadLs. In general, the reactivity of SS14 IRS with Nichols ECLs mirrored that observed with Nichols IRS, with ECL2 and ECL4 of TP0856 and TP0858 again the antigenic standouts (**Fig 6B**). TP0865 ECL3, on the other hand, demonstrated no ELISA reactivity with SS14 IRS. Immunoblot and ELISA with an SS14 TP0865 ECL3 construct revealed that this result was due to the lack of Abs, not sequence variation (**S6 Fig**).

**Fig 6.**
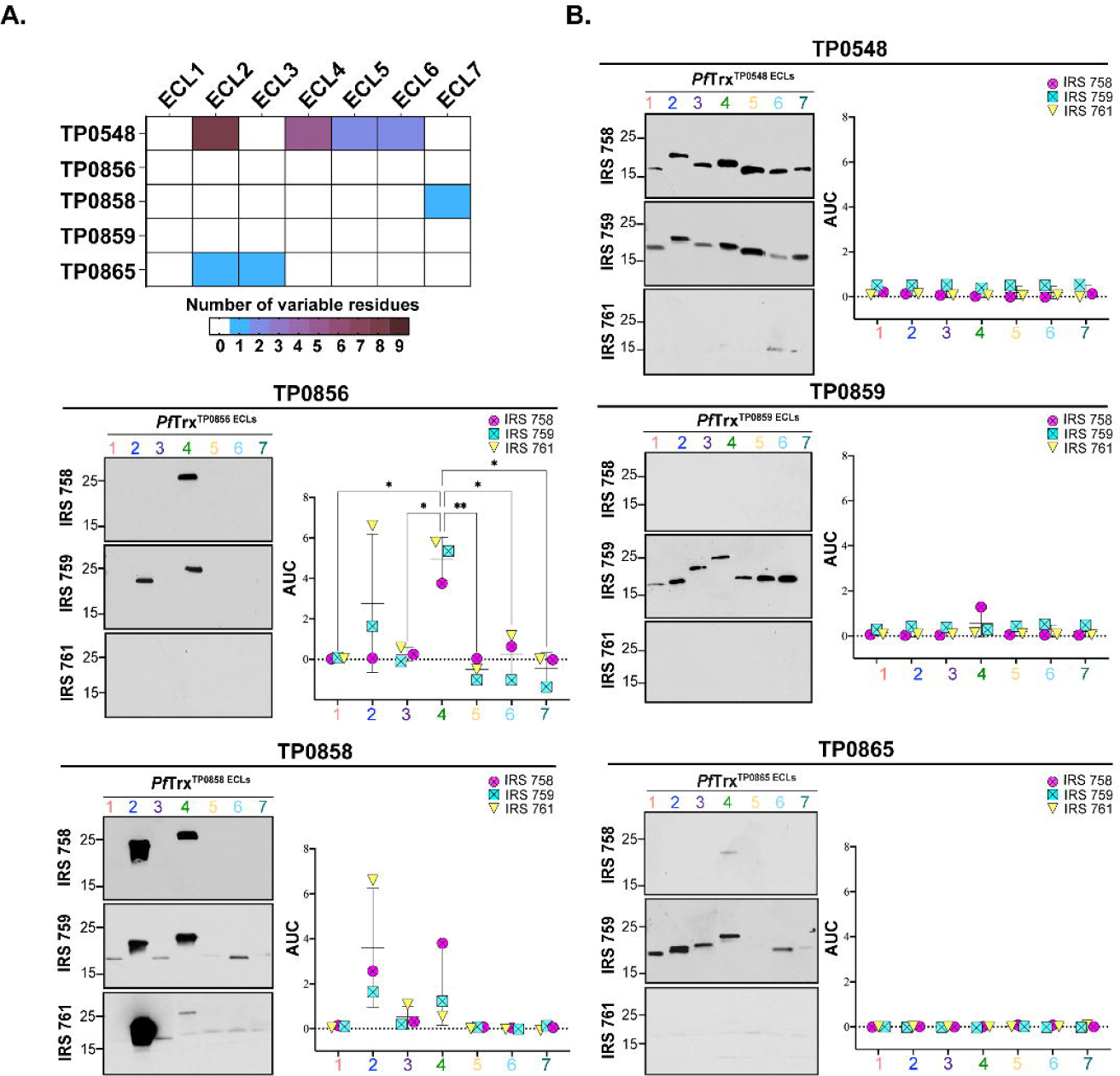
Comparative sequence analysis of the Nichols and SS14 FadLs and reactivity of Nichols FadL ECLs with SS14 IRS. (**A**) Summary chart representing the number of variable residues within Nichols and SS14 FadLs. (**B**) Reactivity by immunoblot (left) and ELISA (right) of Nichols FadL ECLs with SS14 IRS. ELISA reactivity measured as AUC corrected for *Pf*Trx background. *n*L=L3 wells per condition. Data are shown as meanL±LSD. Significant differences (\**p*<0.05 or \*\**p*<0.01) between the means determined by one-way ANOVA with Bonferroni’s correction for multiple comparisons.

### FadL ECL-specific Abs are opsonic for rabbit and murine macrophages

We next sought to determine whether immunization with *Pf*Trx scaffolds displaying immunodominant FadL ECLs would elicit Abs that recognize their native counterparts on *TPA*. We first confirmed the presence of ECL-specific Abs in the rabbit *Pf*Trx^ECL^ antisera by immunoblot and ELISA against the corresponding loops displayed by a heterologous TbpB-LCL scaffold (**Fig 7A** and **7B**)[18, 29]. It was noteworthy that there did not appear to be a strict correlation between the two assays. For example, *Pf*Trx^TP0856/ECL4^ Abs exhibited the strongest ELISA reactivity, yet showed the weakest reactivity by immunoblot. Conversely, *Pf*Trx^TP0856/ECL2^ Abs displayed strong reactivity by immunoblot but the lowest reactivity by ELISA (**Fig 7A** and **7B**). The similar amino acid sequences of the ECL2s and ECL4s in TP0856 and TP0858 (**S7 Fig**) raised the possibility that each ECL might react with the corresponding heterologous antiserum. We investigated this issue using the TbpB-LCL scaffolded ECLs. The ECL2s displayed virtually no cross-reactivity by immunoblot; however, weak cross-reactivity was observed by ELISA (**S7A Fig**). For the ECL4s, weak cross-reactivity was observed by both immunoblot and ELISA (**S7B Fig**). These results indicate that cross-reactivity is not a major confounder for interpreting the results for each ECL antiserum.

**Fig 7.**
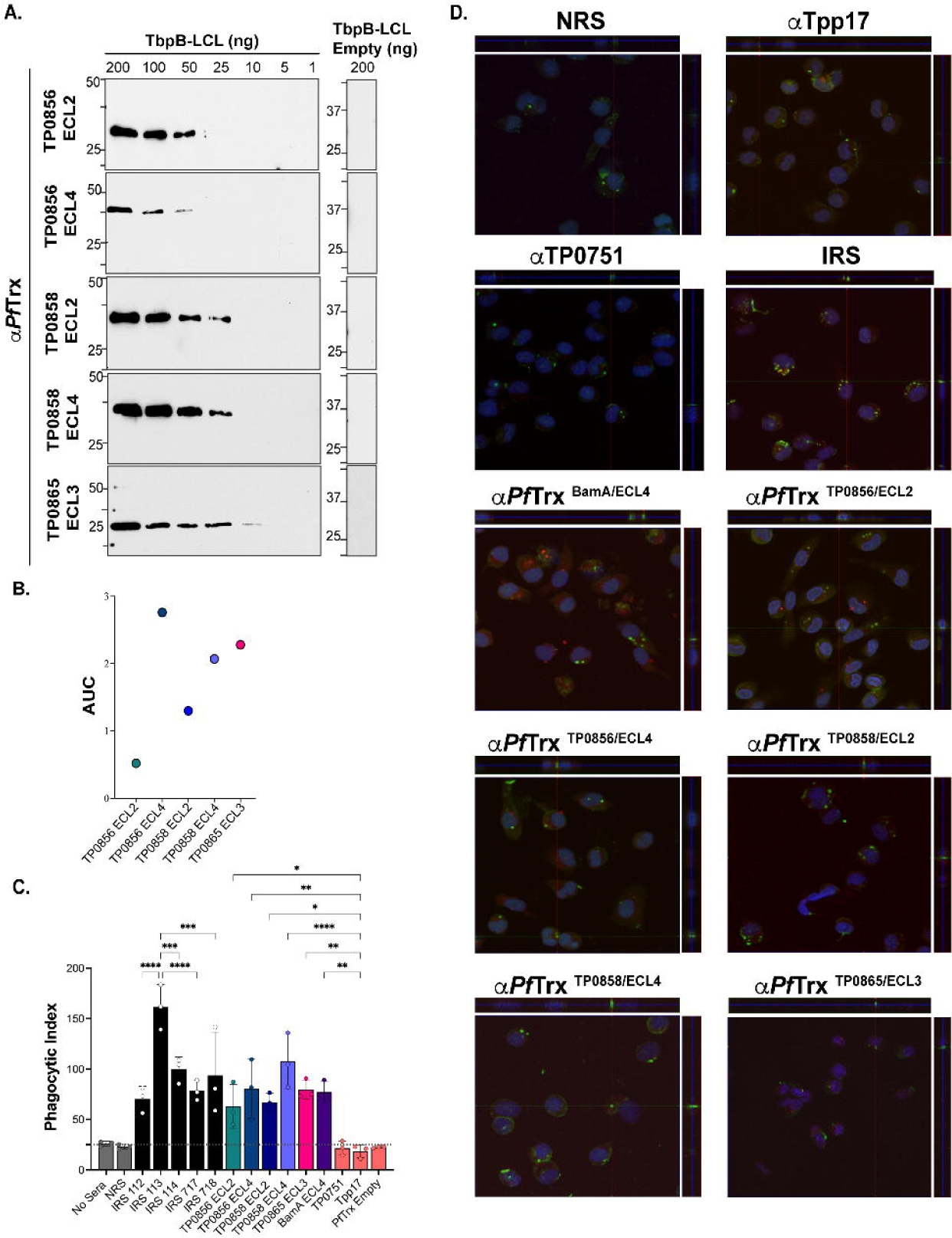
Opsonic activity of rabbit antisera to *Pf*Trx-scaffolded FadL ECLs. (**A**) Immunoblot and (**B**) ELISA (AUC) reactivities of rabbit ECL antisera against the corresponding Tbpb-LCL^ECL^ and Tbpb-LCL^Empty^. (**C**) *TPA* freshly harvested from rabbits was pre-incubated for 2 h with 10% heat-inactivated NRS, IRS, or rabbit antisera to *Pf*Trx ECLs, Tpp17, or TP0751 followed by incubation with rabbit peritoneal macrophages for 4 h at an MOI 10:1. Phagocytic indices were determined from epifluorescence micrographs as described in Methods[18]. Significant differences (\**p*<0.05, \*\**p*<0.01, \*\*\**p*<0.001 or \*\*\*\**p*<0.0001). Bars represent meanL±LSD, *n* = 3 wells per condition. (**D**) Representative confocal micrographs showing composites of 9-12 consecutive Z-stack planes with labeling of *TPA*, plasma membranes, and nuclei shown in green, red and blue, respectively.

For the rabbit opsonophagocytosis assays, sera from the five Nichols immune rabbits and rabbit antiserum against *Pf*Trx-scaffolded ECL4 of BamA/TP0326 (*Pf*Trx^BamA/ECL4^), previously demonstrated to be strongly opsonic[18, 46], served as positive controls; NRS, rabbit α-*Pf*Trx^Empty^, and α-Tpp17 and α-TP0751, previously shown to be non-opsonic[18, 47], were the negative controls. Internalization of spirochetes was assessed using confocal microscopy and quantified by calculating the phagocytic index as described previously[18] and in Methods. Compared to the negative controls, all five IRS exhibited significant opsonic activity, with IRS 113 displaying significantly greater opsonic activity relative to the other four. All five *Pf*Trx FadL ECL antisera demonstrated opsonic activity comparable to IRS; α-*Pf*Trx^TP0858/ECL4^ displayed the most robust opsonic activity (p<0.0001) relative to the negative controls (**Fig 7C** and **7D**).

We recently described an opsonophagocytosis assay employing murine BMDMs to evaluate the opsonic activity of murine monoclonal and polyclonal ECL Abs[18]. As before, we first confirmed the presence of ECL-specific Abs in pooled murine *Pf*Trx^ECL^ antisera by immunoblot and ELISA (**Fig 8A** and **8B**). Immunoblot analysis revealed that three of the five murine ECL antisera (α-*Pf*Trx^TP0856/ECL2^, α-*Pf*Trx^TP0856/ECL4^, and α-*Pf*Trx^TP0858/ECL4^) exhibited comparable sensitivity to their rabbit counterparts, while two (α-TP0858 ECL2 and -TP0865 ECL3) displayed slightly lower reactivity (**Fig 8A**). As with the rabbit ECL antisera, immunoblot and ELISA results obtained with the two assays did not consistently correlate. For example, Abs generated by *Pf*Trx^TP0856/ECL4^ exhibited the weakest ELISA reactivity, despite strong immunoblot reactivity, while *Pf*Trx^TP0858/ECL2^ displayed the strongest ELISA reactivity but low immunoblot reactivity (**Fig 8A** and **8B**). Controls in the murine assay were analogous to the control rabbit sera (described above). Four of the five pooled mouse *Pf*Trx ECL antisera (ECL2s and ECL4s of TP0856 and TP0858) exhibited significant opsonic activity comparable to the pooled mouse syphilitic sera (MSS). Unlike its rabbit counterpart, mouse α-*Pf*Trx^TP0865/ECL3^ was not opsonic (**Fig 8C** and **8D**).

**Fig 8.**
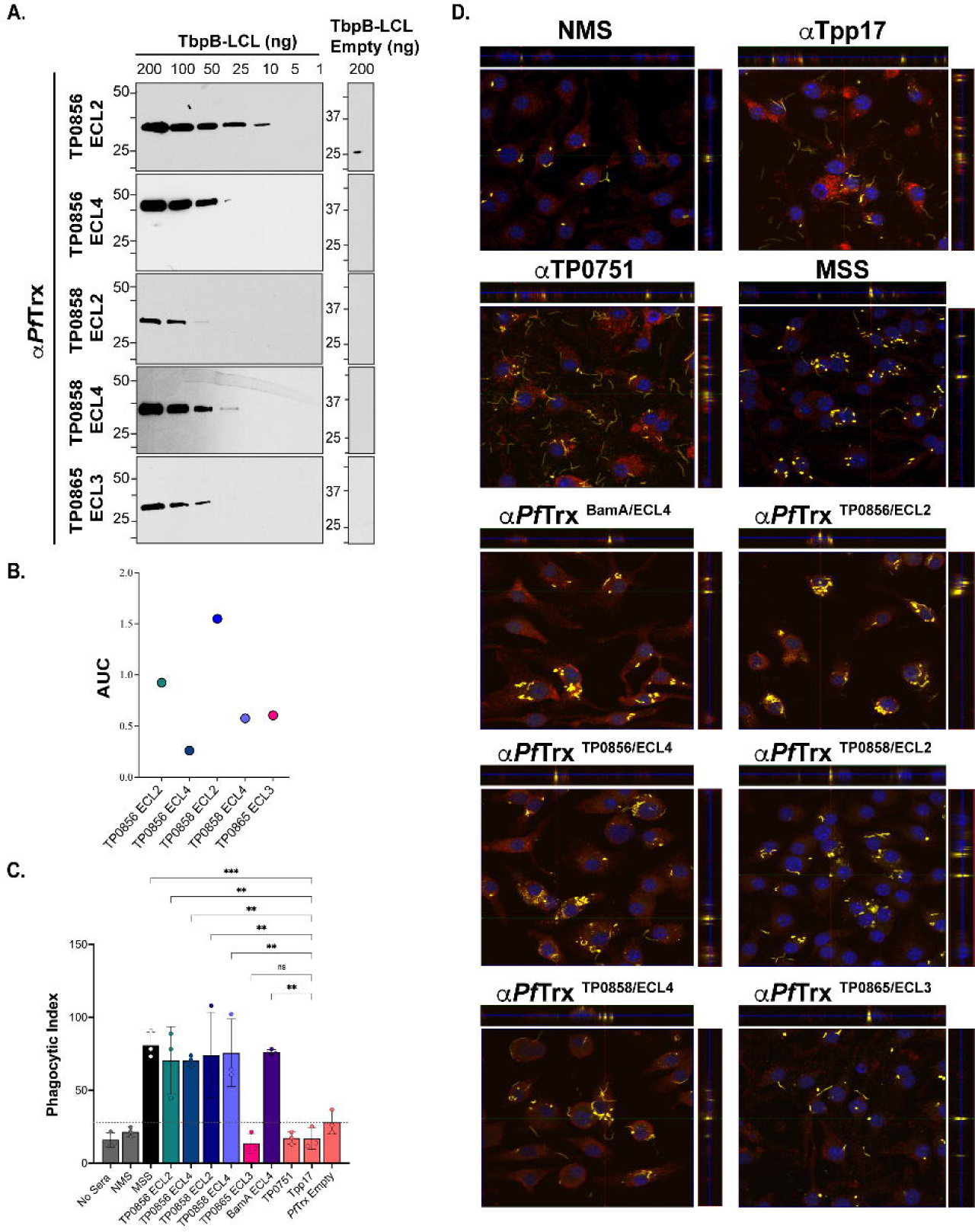
Opsonic activity of mouse antisera to *Pf*Trx-scaffolded FadL ECLs. (**A**) Immunoblot and (**B**) ELISA (AUC) reactivities of sera from mice immunized with *Pf*Trx scaffolded TP0856 ECL2 and ECL4, TP0858 ECL2 and ECL4, and TP0865 ECL3 against graded the corresponding Tbpb-LCL^ECL^ and Tbpb-LCL^Empty^. (**C**) *TPA* freshly harvested from rabbits were pre-incubated for 2 h with 10% heat-inactivated NMS, MSS, and mouse antisera against *Pf*Trx ECLs, Tpp17, and TP0751 followed by incubation with mouse BMDMs for 4 h at an MOI 10:1. Internalization of spirochetes was quantified from epifluorescence micrographs using the phagocytic index. Significant differences (\**p*<0.05 or \*\**p*<0.01). Bars represent meanL±LSD, *n* = 3 wells per condition. (**D**) Representative confocal micrographs showing composites of 9-12 consecutive Z-stack planes with labeling of *TPA*, plasma membranes, and nuclei shown in yellow, red, and blue, respectively.

### Immune sera and ECL-specific Abs exhibit Fc receptor-independent functional activity against *in vitro* cultivated *TPA*

As described in Methods, we modified the recently developed system for continuous *in vitro* propagation of *TPA*[32, 48] to investigate whether heat-inactivated IRS and ECL-specific Abs exert Fc receptor (FcR)-independent functional activity against live *TPA*. Incubation of spirochetes with 10%, 5%, and 1% IRS 112 resulted in a reduction of spirochete numbers below the input level, accompanied by a striking loss of motility and showing severe deterioration of spirochetes (**Fig 9A** and **S1 Movie**), whereas NRS was without effect (**Fig 9A** and **S2 Movie**). Furthermore, unlike NRS, all incubations with IRS contained debris (**S1** and **S2 Movies**). This observation, coupled with the decreased number of spirochetes, points to a bactericidal activity of IRS resulting in spirochete lysis. In our hands, approximately 80% of spirochetes are adherent to the epithelial cells at the time of passage (**S2 Table**). At all three concentrations, IRS also markedly decreased attachment (∼31% attached; *p*<0.0001). In accord with the opsonophagocytosis assays, we saw no effect on growth, motility, or attachment when spirochetes were cultured with α-Tpp17 or α-TP0751 (**Fig 9A**, **S3** and **S4 Movies,** and **S2 Table**). An important question is whether heterologous IRS exerts functional activity in this *in vitro* system. To address this, we compared the impact of incubation with homologous and heterologous IRS on *in vitro*-cultivated Nichols and SS14 *TPA*. Spirochete numbers fell below input levels and motility decreased following incubation of both reference strains with heterologous IRS although the effect was more pronounced with homologous IRS (**Fig 9B** and **9C**).

**Fig 9.**
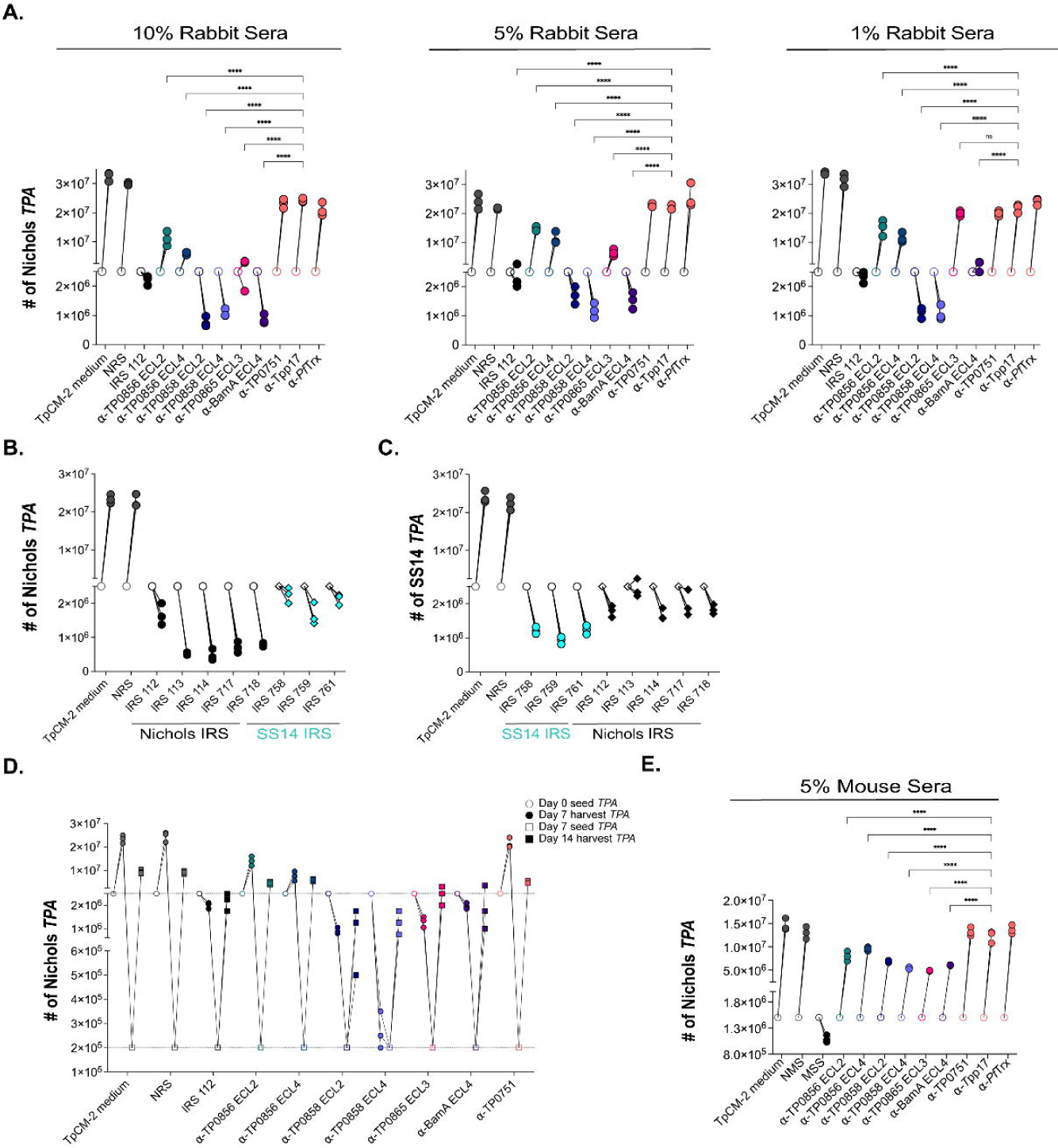
Rabbit and mouse antibodies impact growth of *TPA* Nichols and SS14 during *in vitro* cultivation. (**A**) Enumeration by darkfield microscopy (DFM) at day 7 (solid circles) of spirochetes cultured with 10%, 5%, and 1% concentrations of the indicated rabbit sera, with initial seeding at 2.5 x 10^6^ per well (open circles). (**B**) Nichols *TPA* stain and (**C**) SS14 strain were seeded initially at 2.5 x 10^6^ per well (open shapes) and cultured with Nichols and SS14 IRS (black and cyan, respectively). On day 7 (solid shapes), spirochetes were harvested and enumerated. Homologous IRS and heterologous IRS are depicted as circles and diamonds, respectively. (**D**) Spirochetes harvested on day 7 (solid circles) were transferred to a fresh plate containing Sf1Ep cells and TpCM-2 without rabbit sera. On day 14 (solid squares), spirochetes were harvested and enumerated. (**E**) Enumeration by DFM of spirochetes (initial seeding 1.5 x 10^6^ per well) with 5% mouse antisera targeting FadL ECLs. On day 7, samples were harvested for analysis as described above. Each condition was performed with n=3 replicates. Significant differences (\*\*\*\**p*<0.0001) were determined by two-way ANOVA with Tukey correction for multiple comparisons.

To examine the *in vitro* functional activity of graded concentrations of ECL-specific Abs (**Fig 9A**), we began with rabbit Abs against BamA ECL4, a known target of bactericidal Abs in *E. coli*[49]. In contrast to α-*Pf*Trx^Empty^, incubation of spirochetes with α-*Pf*Trx^BamA/ECL4^ at 10% and 5% resulted in numbers below input levels, whereas growth was static following incubation with 1% α-*Pf*Trx^BamA/ECL4^. All three concentrations resulted in the presence of debris, loss of motility, and a substantial decrease in spirochete attachment (**Fig 9A**, **S2 Table**, and **S5 Movie**). At all three concentrations, incubation with α-*Pf*Trx^TP0858/ECL2^ and α-*Pf*Trx^TP0858/ECL4^ led to reduced spirochete numbers and loss of motility, whereas neither antiserum interfered with attachment. In contrast, only 10% α-*Pf*Trx^TP0865/ECL3^ affected spirochete numbers, motility, and attachment. Significantly, debris consistently were observed with spirochetes incubated with antisera against TP0858 ECLs at all concentrations and 10% α-*Pf*Trx^TP0865/ECL3^. Surprisingly, α-*Pf*Trx^TP0856/ECL2^ and α-*Pf*Trx^TP0856/ECL4^ only modestly affected spirochete growth and had no effect on motility, with only α-*Pf*Trx^TP0856/ECL2^ diminishing attachment (**Fig 9A** and **S2 Table**).

We next sought to determine whether spirochetes could recover from incubation with IRS and α-ECL Abs. In these experiments, we reduced the input organisms into wells without Abs to 2 x 10^5^ spirochetes to compensate for the lower number of treponemes recovered from cultures with IRS and some ECL Abs. While we observed recovery of spirochetes initially cultured with IRS and ECL antisera, none reached counts comparable to those of spirochetes initially incubated with NRS or TpCM-2 medium (**Fig 9D**).

Lastly, we asked whether opsonic murine Abs are functional in the *in vitro* cultivation system. Due to the limited availability of mouse sera, the assay was scaled down and performed at a single concentration (*i.e.*, 5%). At this concentration, MSS reduced total spirochete numbers below the initial seeding amounts and significantly impaired attachment. As expected, NMS and mouse α-Tpp17 and α-TP0751 lacked activity (**Fig 9E**). Interestingly, unlike IRS, MSS did not significantly affect motility and did not result in the presence of debris (**S6-S9 Movies**). In contrast to the rabbit ECL antisera, none of the mouse ECL antisera, including α-*Pf*Trx^BamA/ECL4^, completely inhibited spirochete growth. Notably, the partial inhibition of growth seen with mouse α-*Pf*Trx^TP0856/ECL2^, α-*Pf*Trx^TP0856/ECL4^, and α-*Pf*Trx^TP0865/ECL3^ was comparable to that observed with the corresponding rabbit antisera (**Fig 9E**). Unexpectedly, none of the mouse ECL antisera, including α-*Pf*Trx^BamA/ECL4^ (**S10 Movie**), affected motility or resulted in debris, while all significantly affected attachment to varying degrees (**S3 Table**).

### Transcriptional analysis confirms expression of OMP targets *in vivo* and *in vitro*

Interpretation of the functional activity of ECL Abs requires knowledge of the expression levels of the corresponding OMPs. We utilized publicly available RNAseq data from De Lay *et al*.[50] to compare OMP transcript levels in spirochetes harvested from rabbits and during *in vitro* cultivation. As shown in Fig 10, transcripts for all OMPs studied herein were detected under both conditions. Interestingly, *tp0326*, whose corresponding protein (BamA) was strongly targeted by ECL4 Abs in our opsonophagocytosis and *in vitro* assays, was expressed at relatively low levels *in vivo* and *in vitro*. Several other OMP genes were expressed at comparably low levels under both conditions, among them *tp0865* whose corresponding protein also was well targeted by ECL-specific Abs (**Fig 5** and **Fig 10**). *tp0967*, *tp0733*, and *tp0859* were expressed at higher levels *in vivo* and *in vitro*, suggesting that their poor immunogenicity cannot be attributed solely to poor expression. *tp0856* and *tp0858* had unusual transcriptional profiles. *In vitro*, *tp0856* was expressed at levels comparable to other OMP genes but displayed markedly higher expression *in vivo*. *tp0858* exhibited significantly higher expression *in vitro* compared to all the OMP genes, with a dramatic increase *in vivo*. The transcript levels for the lipoprotein-encoding genes *tp0751* and *tp0435* represented interesting comparators to the OMPs. Both demonstrated higher overall expression levels than many OMPs, with *tp0435* being the only gene with significantly higher expression *in vitro*. The relatively high transcript levels of *tp0751* were unexpected, given the extremely low abundance of the corresponding lipoprotein[47]. Using mass spectrometry (MS)-based proteomics analysis of *TPA* cultivated *in vitro*, Houston *et al*.[51] demonstrated that all the OMPs described herein are expressed by *TPA* at detectable levels. Among them, TP0858 ranked among the top 50 most abundant proteins detected *in vitro*[51]. Additionally, all OMPs, with exception of TP0698 were also detected in *TPA in vivo* from harvested rabbit testes[52–54].

**Fig 10.**
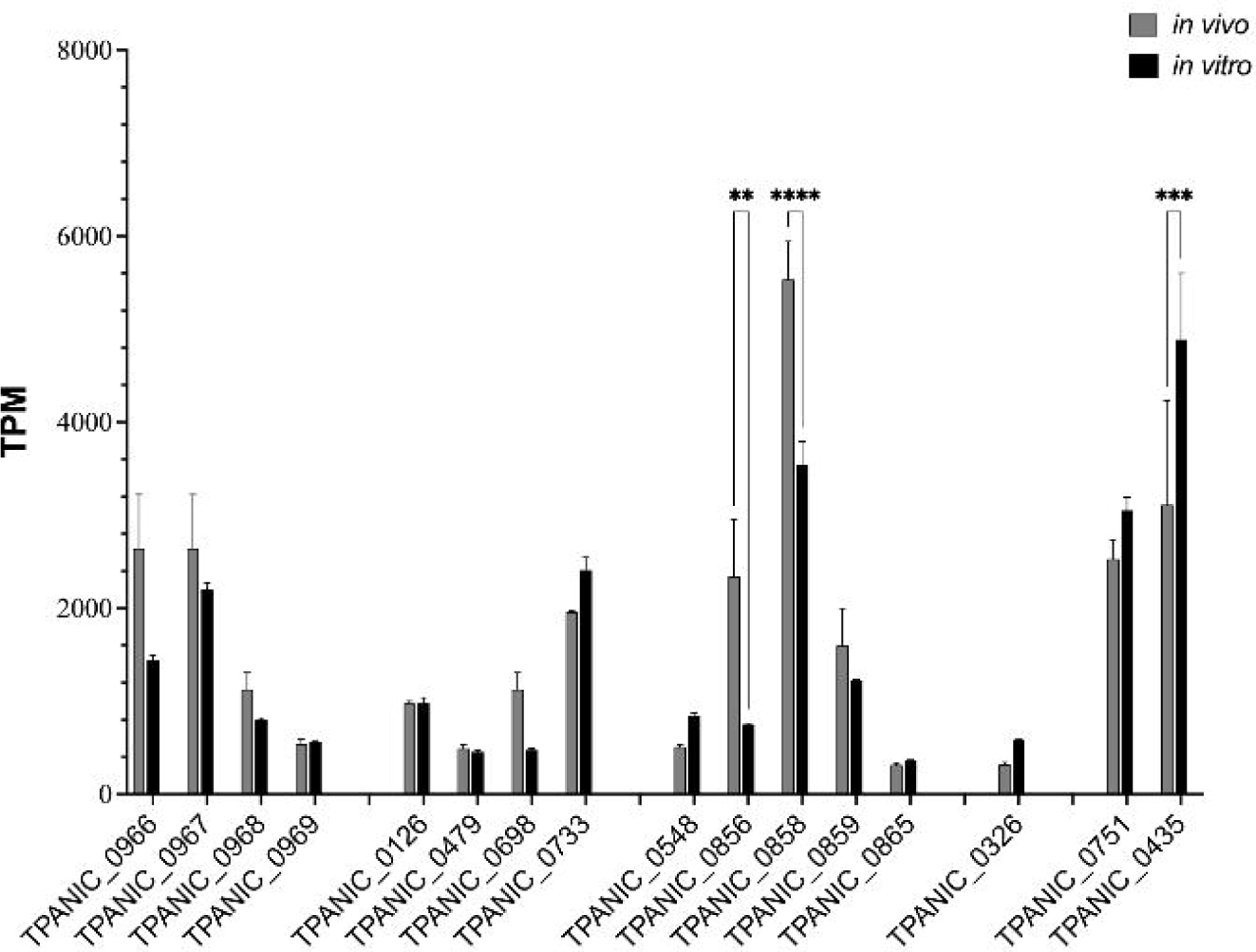
Transcriptional analysis of *TPA* OMP genes. *In vivo* (grey) and *in vitro* (black) expression of *TPA* OMP genes represented as Transcripts per Kilobase Million (TPM) extracted from the RNAseq datasets published by De Lay *et al*.[50].

## Discussion

The alarming global resurgence of syphilis in the twenty first century[1–3] has created an urgent need for a vaccine with worldwide efficacy[4, 5]. A crucial first step for syphilis vaccine development is the identification of *TPA* surface antigens targeted by the functional Abs in immune sera[10–13]. Our strategy for mining IRS for surface-directed Abs was guided by our understanding of the molecular architecture of the *TPA* outer membrane and the structural biology of its repertoire of β-barrel forming OMPs[20–25]. The ‘learning from nature’ variant of rational vaccine design[55] we devised, employing ECLs scaffolded by *Pf*Trx, enabled us to sidestep cumbersome experimentation with full-length OMPs and focus instead on their Ab accessible regions. The success of the approach hinged on the structural models used to define ECL boundaries. Three lines of evidence supported the accuracy of the models. One was the agreement between trRosetta and AlphaFold3 for both 8SβBs and FadLs, alongside the similarity of predicted trRosetta OMFs to crystal structures of OMF orthologs of gram-negative bacteria. Because BCEs are solvent-exposed[56], the location of most predicted BCEs in ECLs provided additional bioinformatic confirmation. Binding of Abs to the surface of motile treponemes, observed in two different assays with rabbit and mouse antisera, provided definitive evidence that the antigenic determinants presented by the scaffolds were extracellular.

ECLs can adopt stable conformations due to interactions with the barrel, with each other[57], or fixed structural elements within the loop[58], while others are mobile and flexible[59–61]. Structural characterization of ECL-Ab complexes reveals that even mobile ECLs adopt specific conformations when bound by bactericidal Abs[62]. Through the use of scaffolds, we discovered that *TPA* ECLs possess a hitherto unsuspected degree of antigenic complexity ostensibly reflecting underlying structural diversity. Instances where ECLs were detected by immunoblot but not ELISA likely indicate linear epitopes that are inaccessible[63] or masked when ECLs are presented in a native-like state. The implications of this observation for disease pathogenesis are rather intriguing. Production of ECL Abs that cannot ‘find’ their linear targets may be a novel manifestation of *TPA*’s capacity for Ab-evasiveness, a virulence trait we have designated ‘stealth pathogenicity[22]. On the other hand, we know from previous work with BamA that some linear epitopes are Ab accessible on scaffolded ECLs as well as live *TPA*[18]. Conversely, reactivity observed by ELISA but not immunoblot presumably results from discontinuous epitopes reproduced when ECLs are tethered. This is an important observation from a vaccine standpoint given the body of evidence that Abs elicited with unfolded OMPs yield an inferior level of protection[64]. Importantly, while certain ECLs, particularly within the FadL family, exhibited robust Ab reactivity in line with predicted BCEs, others, most notably within the OMF and 8SβB families, exhibited poor immunogenicity despite equally strong predictions. Transcriptional and proteomics data indicated that these discordances cannot be attributed to differences in expression. Two explanations, not mutually exclusive, can, therefore, be envisioned. An obvious one is the many limitations known to be associated with BCE predictive algorithms[65]. Another is in line with the presumed poor immunogenicity of the syphilis spirochete’s rare OMPs – the central tenet of the stealth pathogenicity concept[23, 66–68]. To escape immune pressure on functionally critical ECLs, *TPA* may have evolved OMPs whose ECL epitopes ‘slip past’ the host’s Ab generation machinery.

Determination of an *in vitro* correlate of protection as an objective, quantitative criterion for a protective immune response is a prerequisite for the development of a vaccination strategy[69]. Strictly speaking, a true correlate of protection for syphilis does not yet exist. However, *in vivo* evidence from the rabbit model for macrophage-mediated clearance of *TPA*[10] has led to the widely accepted belief that Abs that promote opsonophagocytosis of *TPA* can be considered a surrogate for a protective response[10, 14]. Studies conducted herein with sera from five immune rabbits demonstrated levels of *TPA* internalization greatly surpassing those observed with Abs against the periplasmic controls, Tpp17 and TP0751[42, 47]. It is interesting to note that the five immune sera from outbred rabbits exhibited a broad spectrum of reactivity to our panel of scaffolded ECLs, and that the IRS with the weakest responses overall (IRS 113) displayed the highest level of *TPA* internalization. Collectively, these results point to the protective capacity of different combinations of ECL Abs, and they suggest that examination of ECLs from members of the *TPA* OMPeome not included in the panel is warranted in the effort to create an optimally efficacious ECL vaccine cocktail.

Opsonophagocytosis of *TPA* is slow, inefficient, and incomplete[12, 70, 71]. These observations reflect not just the spirochete’s low density of OMPs but also their poor mobility[23, 72], a physical property that impedes the clustering required for FcR signaling[73]. They also raise the question of whether the infected human host must deploy additional Ab-mediated functions to effect clearance of spirochetes. Years ago, Nelson and Mayer[74] and Bishop and Miller[75] demonstrated *in vitro* complement-dependent killing of *TPA* by syphilitic sera. Azadegan *et al.*[76]showed that depletion of complement in hamsters by administration of cobra venom factor accelerated lesion development following intradermal challenge and prevented protection following passive protection with immune hamster serum. The recent breakthrough in long-term *in vitro* cultivation of *TPA*[32] provided a vehicle to assess whether surface-directed Abs in IRS exert FcR-independent activity against the syphilis spirochete. The detrimental impact of IRS on *TPA* growth and motility, together with the presence of debris not observed with NRS or periplasmic controls, suggested that IRS Abs can exert bactericidal activity. Nevertheless, the ability of spirochetes to recover once immune pressure was relieved points to the presence of a subpopulation of spirochetes capable of surviving the IRS Ab onslaught. This inference aligns with labeling experiments showing extreme variability in the degree of surface Ab binding by IRS within *TPA* populations[20] the survival of subpopulations of spirochetes during opsonophagocytosis experiments[12, 71], and passive-transfer experiments demonstrating the need for continuous administration of IRS to prevent lesion development.[77, 78] We also observed growth inhibition and lack of motility of Nichols and SS14 *TPA* cultured with homologous and heterologous IRS strains *in vitro*. These findings imply that Abs directed against conserved ECL epitopes may result in cross-immunity. On the other hand, homologous IRS caused a more pronounced effect on *TPA* viability than heterologous IRS, supporting the importance of Abs against variable surface epitopes for full protection. That Abs in IRS exert FcR-dependent and -independent activities clearly works to the advantage of the host. Organisms immobilized or killed by IRS would be ‘sitting ducks’ for tissue macrophages. Cellular immunity also plays an important role in this scenario since macrophages require activation by IFN-γ produced by locally infiltrating T cells to internalize Ab-opsonized treponemes[13]. In addition to affecting growth and motility, IRS markedly impaired *TPA* attachment to rabbit epithelial cells. The ability of IRS to prevent cytoadherence of *TPA* to multiple cell types is well described[79–81]. Abs against TP0751 did not interfere with attachment, supporting previous data from our group that this protein, rather than being a surface adhesin/protease[82, 83], is a low abundance, periplasmic lipoprotein possibly involved in heme acquisition[47].

The ‘learning from nature’ paradigm for vaccine design rests on the premise that immunization with immunogenic surface molecules identified in an immune serum will, if properly formulated, yield functional Abs[55]. This premise clearly was fulfilled for all five scaffolded, immunodominant FadL ECLs mined from IRS. Until recently[18], opsonophagocytosis assays were conducted with Abs to full-length proteins or protein domains[46, 84–87]; positive results with these antigens left open the question of the precise surface location of the opsonic epitopes. Use of ECLs resolves this issue at a topological level though the specific residues involved in Ab binding still needs to be determined structurally. Opsonophagocytosis requires that Abs bind to the bacterial surface for recognition by FcRs; the Abs, however, are not the effectors. Studies with the *in vitro* cultivation system revealed that Abs targeting specific ECLs can be true effectors, interfering with the functions of individual OMPs to cause severe, even fatal, physiologic perturbations reflected by loss of motility and viability.

BamA is a central component of a molecular machine that cycles between open and closed states to insert newly synthesized OMPs into the OM bilayer[49, 88]. ECL4 is part of a multi-loop dome that prevents egress of the OMP substrates to the external milieu^88^. Presumably, Ab binding to ECL4 prevents movements within the dome needed to accommodate cycling of the BamA β-barrel, inflicting a fatal lesion by impairing OM biogenesis. Growth inhibition and killing by anti-FadL ECL Abs undoubtedly reflects interference with uptake of essential small molecules, though the mechanism is unclear in light of current thinking about how FadLs capture hydrophobic substrates and direct them into and through the β-barrel[36]. We observed an intriguing dichotomy with Abs to ECLs 2 and 4 of TP0856 and TP0858. Abs to the TP0858 ECLs had a dramatic effect on *TPA* growth and survival while the effects of Abs to the corresponding loops of TP0856 were comparatively weak. Given the similar opsonophagocytosis results with these same Abs, differences in Ab binding seem implausible; more likely is that TP0856 is physiologically redundant within the *in vitro* environment. Abs against two ECLs, ECL4 of BamA and ECL2 of TP0856, had a pronounced effect on cellular attachment, though in the context of markedly different effects on growth and motility. It seems reasonable to conjecture that the anti-adhesive effect of the BamA ECL4 Abs was the result of a broad derangement of the *TPA* surface, while the TP0856 Abs ostensibly interfered with a *bona fide* ECL-dependent adhesive function. This supposition is in line with numerous examples of bacterial OMPs involved in maintaining cellular homeostasis whose ECLs have a virulence-related function as adhesins[89–92]. The *in vitro* cultivation system promises to be an important addition to the syphilologist’s toolkit for dissecting the cytadhesive properties of *TPA* OMPs - an area of investigation at the nexus of vaccine development and syphilis pathogenesis.

The rabbit has been the animal model of choice for basic syphilis research for decades[6–8]. Our studies deciphering ECL Ab responses in animals with proven immunity to intradermal inoculation and then improving upon them by artificial immunization further demonstrate the model’s utility. Nevertheless, the outbred nature of the rabbit, the skyrocketing costs for purchase and maintenance, and the limited commercial availability of rabbit-specific reagents impose serious constraints at a time of great urgency for identification, refinement, and validation of protective targets. Historically, the lack of skin lesion development, the large inoculum required for infection, and the delayed time course for spirochete clearance have discouraged use of the mouse model[15, 16, 19]. Moreover, whether mice develop protective immunity has not yet been established. While the immunobiology of syphilis in the mouse may be less than optimal for pathogenesis studies, the evidence in hand points to the mouse as the obvious animal model for expediting vaccine research. *TPA*-infected mice generate Abs that strongly promote phagocytosis of spirochetes by BMDMs[18], as well as Abs that inhibit *TPA* growth *in vitro*. From these results, we can surmise that *TPA-*infected mice, like rabbits, develop Abs directed against ECLs and that comparison of the two responses could be highly informative. Overall, however, the murine responses following immunization with scaffolded ECLs were less robust than those of rabbits; this was particularly evident from the *in vitro* cultivation experiments. From one perspective, these differences are advantageous since they can be exploited to pinpoint ECL epitopes most important for protective Abs. On the other hand, strategies to improve them clearly will need to be devised before the mouse can take its place as a reliable screening tool. Despite lingering questions and historical prejudices, the mouse model brings to syphilis vaccinology unparalleled benefits, including cost-effectiveness, access to a vast array of reagents, and a wealth of inbred strains with precisely defined genetic backgrounds and fully characterized immunologic phenotypes.

## Materials and Methods

### Ethics statement

Animal experimentation was conducted following the *Guide for the Care and Use of Laboratory Animals* (8th Edition) in accordance with protocols reviewed and approved by the UConn Health Institutional Animal Care and Use Committee (AP-200351-0124, AP-200362-0124, AP-201085-1226, and AP-201086-1226) under the auspices of Public Health Service assurance number A3471-01 (D16-00295).

### OMP Modeling

Three-dimensional models for the OMFs (TP0966, TP0967, TP0968, and TP0969), 8SβBs (TP0126, TP0479, TP0698, TP0733), and FadLs (TP0548, TP0856, TP0858, TP0859, and TP0865) were retrieved from pre-existing models generated from Hawley *et al*.[24]. For all three families, structural models and ECL boundaries were re-examined using AlphaFold3[34] (https://golgi.sandbox.google.com/).

### B cell epitope analysis

Linear and conformational B cell epitopes (BCEs) were predicted from the trRosetta 3D models using DiscoTope 2.0[40] and ElliPro[39] (**S1Table**). We used a threshold ≥0.8 for conformational BCE predictions by ElliPro and default settings for DiscoTope.

### Propagation of *TPA* and generation of immune rabbit sera

The *TPA* Nichols and SS14 reference strains (SS14 was generously provided by Dr. Steven Norris, McGovern Medical School, University of Texas Health Science Center at Houston) were propagated by intratesticular inoculation of adult male New Zealand White (NZW) rabbits as previously described[13, 20]. Immune rabbits were generated by inoculation of rapid plasma reagin-nonreactive adult NZW rabbits in each testis with 1 x 10^7^ treponemes in 500 μL CMRL containing 20% NRS. The immune status of each rabbit was confirmed sixty days post-inoculation by intradermal challenge with 1 x 10^3^ freshly extracted *TPA* (Nichols or SS14) at each of eight sites on their shaved backs. Immune sera were collected at monthly intervals thereafter.

### Generation of mouse syphilitic sera

Male and female six- to eight-week-old C3H/HeJ mice were inoculated intradermally, intraperitoneally, intrarectally, and intra-genitally with a total of 1 x 10^8^ total organisms per animal as previously described[17, 18]. Mice were sacrificed on day 84 post-inoculation and exsanguinated to create a pool of MSS.

### Cloning ECLs into *Pf*Trx and TbpB-LCL scaffolds

A codon-optimized version of *Pyrococcus furiosus* thioredoxin (*Pf*Trx)[27] with *TPA* BamA ECL4 inserted between amino acid residues 26 and 27 of the native *Pf*Trx and a C-terminal Avi-Tag (GLNDIFEAQKIEWHE) was synthesized by Genewiz. The resulting construct (*Pf*Trx^BamA/ECL4^) was PCR-amplified and cloned into NdeI-XhoI digested pET28a by In-Fusion cloning. To generate *Pf*Trx^Empty^, *Pf*Trx^BamA/ECL4^ was digested with BamHI to remove the ECL4-encoding DNA and then self-ligated. Supporting Table 1 contains the primers and sequences used to generate amplicons encoding ECLs other than BamA ECL4 for display by *Pf*Trx scaffolds (see below). *Pf*Trx scaffolds displaying ECLs shorter than 30 amino acids were generated by inverse PCR of pET28a*^Pf^*^Trx^ using primers containing the corresponding ECL sequences followed by InFusion cloning. *Pf*Trx constructs containing ECLs longer than 30 amino acids were generated by PCR-amplifying the loops from codon-optimized synthetic genes followed by insertion into BamHI-digested pET28a*^Pf^*^Trx^ by InFusion cloning. *Pf*Trx ECLs used for antigenic analyses (see below) were biotinylated during expression in *E. coli* BL21 (DE3) transformed with BirA (BPS Bioscience, San Diego, CA)[27].

DNAs encoding transferrin-binding protein B loopless C-lobe scaffold (TbpB-LCL) derived from *Neisseria meningitidis* TbpB[18, 29] and TbpB-LCL displaying *TPA* ECLs (**S1 Table**) were synthesized by Azenta Life Sciences (Burlington, MA) and cloned into pRB1B by In-fusion cloning as previously described[18, 27]. Plasmid inserts were confirmed by Sanger sequencing and then transformed into *E. coli* BL21-Gold (DE3) (Agilent, Santa Clara, CA) for overexpression. All constructs were purified over Ni-NTA resin (Qiagen, Germantown, MD) followed by size exclusion chromatography as previously described[27].

### Immunoblot analysis of IRS with *TPA* lysates

Nichols *TPA* lysates (5 x 10^7^ spirochetes per lane) were resolved by SDS-PAGE using a 4-20% gradient Any kD Mini-Protean TGX gels (Bio-Rad) and transferred to 0.45 nm nitrocellulose membranes (Bio-Rad, Hercules, CA). The membranes were blocked for 1 h with PBS containing 5% nonfat dry milk and 0.1% Tween 20 and probed overnight (ON) at 4°C with either Nichols or SS14 immune rabbit serum (IRS) (both at 1:1,000 dilutions) from individual rabbits. After washing with PBS containing 0.05% Tween 20 (PBST), the membranes were incubated for 1h at RT with HRP-conjugated goat anti-rabbit IgG or (1:30,000). Following further washes with PBST, the immunoblots were developed on a single film using the SuperSignal West Pico chemiluminescent substrate (ThermoFisher Scientific, Inc., Waltham, MA).

### Reactivity of rabbit and mouse syphilitic sera with *Pf*Trx-scaffolded ECLs

#### Immunoblot

400 ng of *Pf*Trx^Empty^, *Pf*Trx-scaffolded ECLs, and 20 ng of Tpp17 were incubated with 1:250 dilutions of Nichols or SS14 IRS or pooled Nichols MSS followed by HRP-conjugated goat anti-rabbit IgG or goat anti-mouse Ig (1:30,000) as described above.

#### ELISA

Clear Flat-Bottom Immuno Nonsterile 96-well plates (ThermoFisher Scientific, Inc.) were coated with streptavidin (SP; ThermoFisher Scientific, Inc.) diluted in 0.1M sodium bicarbonate (pH 8.5) at 200 ng/well and incubated ON at 4°C. After washing with 0.1% PBST, plates were blocked in PBS buffer containing 15% goat serum, 0.5% Tween 20, and 0.05% sodium azide (blocking buffer) for 1 h at RT. Biotinylated ECL scaffolded proteins were added at 200 ng/well in blocking buffer followed by incubation for 1 h at RT. After washing, either Nichols IRS, SS14 IRS, or MSS was added in 2-fold serial dilutions (1:20 starting dilution) in PBS with 1% bovine serum albumin (BSA) for 1 h incubation at RT. HRP-conjugated goat anti-rabbit IgG or goat anti-mouse Ig (1:10,000) then was added, followed by incubation for 1 h at RT. Plates were washed and developed with TMB single solution (ThermoFisher Scientific, Inc.). Reactions were stopped with 0.3M HCl. Area under the curve (AUC) for each scaffolded ECL were calculated following subtraction of the AUC for *Pf*Trx^Empty^.

### Sequence alignment of Nichols and SS14 FadLs

Protein sequences for full length FadL orthologs or selected ECLs from the *TPA* Nichols (CP004010.2) and SS14 (CP004011.1) reference genomes were aligned using Clustal Omega[93].

### Immunization of rabbits and mice with *Pf*Trx-ECLs

Adult male NZW rabbits were primed with a total of 200 μg of *Pf*Trx-scaffolded ECL in 500 μl of PBS-TiterMax (1:1, vol/vol) administered as four subcutaneous injections and two intramuscular injections with 100 μL and 50 μL, respectively. Rabbits were boosted at 3, 6, and 9 weeks with the same volumes and amounts of protein in PBS/TiterMax (1:1, vol/vol) and exsanguinated 12 weeks post-immunization. Six- to eight-week-old C3H/HeJ mice (Jackson Laboratory) were primed by intradermal injections with 100 μl Freund’s Complete Adjuvant (1:1, v/v) containing 20 μg of ECL scaffolded proteins described above. Mice were boosted at 3, 5, and 7 weeks with the same volumes and amounts of protein in Freund’s Incomplete Adjuvant (1:1, v/v) and exsanguinated 9 weeks post-immunization. Sera from rabbits and pooled sera from mice were heat-inactivated, and then used in immunologic assays.

### Characterization of ECL-specific Abs in *Pf*Trx ECL antisera

#### Immunoblotting

ECL-specific reactivity of rabbit and mouse *Pf*Trx-ECL antisera was determined using TbpB-LCL from *Neisseria meningitis* as a second ECL scaffold as previously described[18]. Graded amounts of the corresponding TbpB-LCL-ECL (200 to 1 ng) were resolved by SDS-PAGE using AnykD Mini-Protean TGX gels, transferred to nitrocellulose, and probed ON at 4°C with 1:1000 dilutions of rabbit or mouse *Pf*Trx-ECL antisera. After washing with PBST, the membranes were incubated for 1 h at RT with HRP-conjugated goat anti-rabbit IgG or goat anti-mouse Ig (1:30,000) as previously described[18]. 200 ng of TbpB-LCL^Empty^ was used as a negative control.

#### ELISA - rabbit antisera

Clear Flat-Bottom Immuno Nonsterile 96-well plates (ThermoFisher Scientific, Inc.) were coated with 6x-His tag monoclonal antibody (HIS.H8) (ThermoFisher Scientific, Inc.) diluted in 0.1M sodium bicarbonate at 200 ng/well and incubated ON at 4°C. All subsequent steps were performed as described above using 200 ng/well of TbpB-LCL-scaffolded ECL and a 2-fold serial dilution of *Pf*Trx^ECL^ antisera. The AUC for each scaffolded ECL was calculated following subtraction of the AUC for TbpB-LCL^Empty^.

#### ELISA - mouse antisera

Mouse *Pf*Trx^ECL^ antisera was absorbed against TbpB-LCL^Empty^ using Dynabeads™ (CAT# 10103D, 10104D) according to the His-Tag Isolation & Pulldown protocol from Invitrogen. ECL-specific Abs were then assessed using Clear Flat-Bottom Immuno Nonsterile 96-well plates (ThermoFisher Scientific, Inc.) coated at 200 ng/well with TbpB-LCL-scaffolded ECLs in PBS. Washes and blocking were performed as described above following 2 h of incubation at RT. After blocking, the corresponding absorbed *Pf*Trx^ECL^ antisera was added at 2-fold serial dilutions (starting at 1:20) in PBS with 1% BSA for a 1 h incubation at RT. HRP-conjugated goat anti-mouse Ig (1:10,000) then was added, followed by incubation for 1 h at RT. The AUC for each scaffolded ECL was calculated following subtraction of the AUC for TbpB-LCL^Empty^.

### Opsonophagocytosis assays

#### Generation of macrophages

Rabbit peritoneal macrophages were generated using 10% protease peptone and isolated using ice-cold PBS EDTA as previously described[18, 47]. The macrophages were plated at a final concentration of 1 x 10^5^ cells/well in 8-well BioCoat Poly-D-Lysine glass culture chamber slides (Corning, Corning, NY) and incubated at 37°C for 2 h. Nonadherent cells were removed by washing the monolayers twice with DMEM prior to the addition of *TPA*. Murine C3H/HeJ bone-marrow-derived macrophages (BMDM) were generated as previously described[17, 18], plated at a final concentration of 1 x 10^5^ cells per well in Millicell EZ 8-well chamber slides (Sigma-Aldrich, St. Louis, MO), and incubated ON at 37°C. The following day, the medium was replaced with fresh Dulbecco’s Modified Eagle Medium (DMEM) supplemented with 10% FBS prior to the addition of *TPA*.

#### Opsonophagocytosis

Freshly harvested *TPA* were diluted to 1 x 10^8^ per ml in DMEM or DMEM supplemented with 1:10 dilutions of normal mouse or rabbit serum, mouse or rabbit syphilitic sera, or mouse or rabbit antisera directed against *Pf*Trx^TP0856/ECL2^, *Pf*Trx ^TP0856/ECL4^, *Pf*Trx ^TP0858/ECL2^, *Pf*Trx ^TP0858/ECL4^, *Pf*Trx ^TP0865/ECL3^, *Pf*Trx^Empty^. Negative controls included rabbit and mouse α-Tpp17 and α-TP0751 sera[42, 47]. Each stimulation condition was performed in triplicate. *TPA* was pre-incubated at RT for 2 h without or with sera followed by incubation for 4 h at 37°C with macrophages (plated as described above) at MOIs of 10:1.

#### Determination of spirochete uptake

Supernatants were removed, and rabbit peritoneal macrophages were fixed and permeabilized with 2% paraformaldehyde and 0.01% Triton X-100 for 10 mins at RT. Each well was rinsed with PBS and blocked with CMRL 10% normal goat sera (NGS) for 1 h at RT, and then incubated with MSS generated above (1:25) in CMRL 10% NGS ON at 4°C. After four successive washes with PBST, cells were blocked with CMRL 10% NGS for 1 h at RT, then incubated with α-mouse IgG AF488 (1:500) for 1 h at RT, followed by Cholera Toxin AF647 (1:500) for 30 min and DAPI (1:1000) for 10 min. After staining for *TPA*, the cells then were washed thoroughly three times with PBST, rinsed with deionized (DI) water to remove salt, and allowed to air dry. Finally, Vectashield^®^ (Vector Laboratories, Inc., Newark, CA) was added, and samples were sealed with a coverslip. Internalization of *TPA* was assessed in a blinded fashion by acquiring images of at least 100 macrophages per well on an epifluorescence Olympus BX-41 microscope[18]; images were processed with VisiView (version 5.0.0.7; Visitron Systems GmbH, Puchheim, Germany). The phagocytic index was calculated by dividing the number of internalized spirochetes by the total number of cells imaged and multiplying by 100[18]. Confocal images were acquired using Zeiss 880 and processed using ZEN3.5 Blue. For IFA of murine BMDMs, cells were blocked with 5% BSA in PBS for 1 h at RT and then incubated with a commercially available rabbit α-*TPA* (1:100), ON at 4°C. the next day the cells were washed four times with PBST and incubated with a-rabbit IgG Texas Red (1:500) for 1 h at RT, followed by Phalloidin AF488 (1:10), Cholera Toxin AF647 (1:500) for 30 min, and DAPI (1:1000) for 10 min. Internalization of *TPA* was assessed as described above.

### Assessment of functional activity using *in vitro* cultivated *TPA*

Cottontail rabbit epithelial cells (Sf1Ep)[32, 48], generously provided by Drs. Diane Edmonson and Steven Norris (UT Health Science Center at Houston), were seeded at 2 x 10^4^ cells/well in a 24-well culture plate and incubated ON at 37°C. The following day, wells were washed once with *TPA* culture medium 2 (TpCM-2)[32, 48] equilibrated under microaerobic conditions (1.5% O_2_, 3.5% CO_2_, and 95% N_2_) followed by the addition of 2.5 ml of fresh TpCM-2 for a minimum of 3 h under microaerobic conditions. 2.5 x 10^6^ freshly harvested *TPA* were added to each well along with normal sera, or syphilitic sera, or *Pf*Trx ECL antisera (*Pf*Trx ^TP0856/ECL2^, *Pf*Trx ^TP0856/ECL4^, *Pf*Trx ^TP0858/ECL2^, *Pf*Trx ^TP0858/ECL4^, *Pf*Trx ^TP0865/ECL3^). Control antisera included *Pf*Trx^BamA/ECL4^, *Pf*Trx^Empty^, Tpp17 and TP0751. Spirochetes were harvested following incubation for seven days under microaerobic conditions. Supernatants were collected and set aside for subsequent DFM enumeration. Wells were then washed once with 200 μl of trypsin EDTA to remove traces of TpCM-2 media. 200 μl of Trypsin EDTA then was added to each well and incubated at 37°C for 5 min following which *TPA* released from the cells was collected in separate 5 ml conical tubes. The supernatant and cell-associated (*i.e*., trypsinized) fractions were centrifuged at 130 x *g* for 5 min followed by DFM enumeration. Movies following incubations were obtained using OCULAR^TM^ Advanced Scientific Camera Control version 2.0 (64 bit) software with PVCAM version 3.8.0 (Teledyne Photometrics, Tucson, AZ)

To evaluate the viability of spirochetes following incubation with IRS and ECL-specific Abs, 2 x 10^5^ spirochetes per well were passaged to fresh wells containing Sf1Ep cells and fresh TpCM-2 medium for an additional 7 days followed by DFM enumeration. In these experiments, the number of input organisms was reduced to normalize for the lower numbers of treponemes harvested from day 7 cultures containing Abs.

### Comparison of *in vivo* and *in vitro TPA* OMPs transcripts

Previously published[50] raw read sequencing data for *TPA* strain Nichols cultivated *in vitro* and harvested from infected rabbits were downloaded from the NCBI Sequence Read Archive (SRA) database (accession numbers SRR16297052, SRR16297053, SRR16297054, SRR16297055, SRR16297056, SRR16297057, SRR16297058 and SRR16297059). Reads were trimmed using Sickle version 1.3.3 (available from https://github.com/najoshi/sickle)[94] and then mapped using EDGE-pro version 1.1.3[95] using fasta, protein translation table (ptt) and ribosomal/transfer RNA table (rnt) files based on the *TPA* strain Nichols genome (RefSeq: NC_021490.2). Transcripts per kilobase million (TPM) values were calculated as previously described[96] using reads mapped to *TPA* protein coding sequences.

### Statistical analysis

General statistical analysis was conducted using GraphPad Prism 9.5.1 (GraphPad Software, San Diego, CA). The means of the AUC from ELISA dilution curves for the *Pf*Trx-scaffolded ECLs constructs were compared by one-way ANOVA with Bonferroni’s correction for multiple comparisons. One-way ANOVA was used to compare phagocytic indices in rabbits and mice using Newman-Keuls and Bonferroni’s correction for multiple comparisons, respectively. A two-way ANOVA was used to compare *TPA* growth *in vitro* with Tukey correction for multiple comparisons in rabbits and mice assays. Ordinary one-way ANOVA was used to compare attached *TPA* in rabbits and in mice using Bonferroni’s correction for multiple comparisons. Two-way ANOVA was used to compare OMP gene transcripts among each other as well as comparison of *in vivo* and *in vitro* OMP transcripts using Šidák correction for multiple comparisons. For each experiment, the standard error of the mean was calculated with *p*-values <0.5 considered significant.

## Acknowledgments

We thank Dr. Steven J. Norris, McGovern Medical School, University of Texas Health Science Center at Houston, for providing the *TPA* SS14 reference strain. We also thank Drs. Steven J. Norris, Diane G. Edmonson, and Bridget D. DeLay for their invaluable support and guidance in establishing the *TPA in vitro* cultivation system in the UConn Health Spirochete Research Laboratories. We thank Ms. Morgan LeDoyt and Mr. Kemar Edwards for outstanding technical support. Lastly, we extend our gratitude to Drs. Trevor F. Moraes (University of Toronto) and Anthony B. Schryvers (University of Calgary) for their continued support with the use of the TbpB-LCL as a protein scaffold.

## Data Availability Statement

The data that support the findings of this study will be available by contacting the corresponding author directly following the date of publication.

## Supporting Information

**S1 Fig. Schematic depicting trRosetta and AlphaFold3 ECL boundary predictions.** One-dimensional schematic for (**A**) OMFs, (**B**) 8SβBs, and (**C**) FadLs depict trRosetta (teal box) and AlphaFold3 (black box) predicted ECLs. ECL boundaries used to clone ECLs onto the *Pf*Trx scaffold are depicted in boxes using color scheme described in **Figure 1**.

**S2 Fig. Comparison of OMF crystal structures with structures for TP0967 predicted by AlphaFold3 and trRosetta**. OMF crystal structures of *Neisseria gonorrhoeae* MtrE and *E. coli* TolC compared to trRosetta and AlphaFold3 three-dimensional models of OMF TP0967.

**S3 Fig. Three-dimensional models depicting B cell epitope (BCE) predictions for OMFs, 8S**β**Bs and FadLs.** 3D models for (**A**) OMFs, (**B**) 8SβBs, and (**C**) FadLs depicting discontinuous BCE predictions by Disco Tope 2.0 (pink surface) and ElliPro (purple surface).

**S4 Fig. Immunoblot reactivity of Nichols and SS14 IRS with *TPA* Nichols lysates, *Pf*Trx^Empty^, and Tpp17.** (**A**) Immunoblot reactivity of Nichols and SS14 IRS with Nichols lysates. (**B**) Immunoblot reactivity of Nichols and SS14 IRS with *Pf*Trx^Empty^ and Tpp17 proteins.

**S5 Fig. Sequence alignments for FadL orthologs in *TPA* Nichols and SS14 reference strains.** Clustal Omega[93] alignments of the five Nichols FadL orthologs with highlighted variations shown in magenta. Predicted ECLs are indicated using color scheme described in **Figure 1**. Discontinuous BCE predictions by DiscoTope 2.0 and ElliPro are shown in purple boxes along the sequences.

**S6 Fig. Antigenic characterization of SS14 TP0865 ECL3.** (**A**) Immunoblot and ELISA (AUC) reactivity of SS14 TP0865 ECL3 with (**B**) Nichols and (**C**) SS14 IRS.

**S7 Fig. Limited cross-reactivity of ECLs 2 and 4 of TP0856 and TP0858.** (**A**) Alignment of TP0856 and TP0858 ECL2 sequences. Immunoblot and ELISA (bottom left and right, respectively) reactivity of rabbit anti-*Pf*Trx^TP0856/ECL2^ and anti-*Pf*Trx^TP0858/ECL2^ against TbpB-LCL^TP0856/ECL2^ and TbpB-LCL^TP0858/ECL2^. (**B**) Alignment of TP0856 and TP0858 ECL4 sequences. Immunoblot and ELISA (bottom left and right, respectively) of rabbit anti-*Pf*Trx^TP0856/ECL4^ and anti-PfTrx^TP0858/ECL4^ against TbpB-LCL^TP0856/ECL4^ and TbpB-LCL^TP0858/ECL4^.

**S1 Table. ECL sequences and primer pairs used to generate *Pf*Trx-scaffolded ECLs.**

**S2 Table. Attachment of *in vitro*-cultivated *TPA* Nichols following incubation for seven days with IRS and rabbit antisera.**

**S3 Table. Attachment of *in vitro*-cultivated *TPA* Nichols following incubation for seven days with pooled MSS and mouse antisera.**

**S4 Table. Statistical analysis of *in vivo* and *in vitro* transcripts for *TPA* OMP genes.**

**S1 Movie. DFM microscopy of *in vitro* cultivated *TPA* Nichols incubated for seven days with 10% IRS.**

**S2 Movie. DFM microscopy of *in vitro* cultivated *TPA* Nichols incubated for seven days with 10% NRS.**

**S3 Movie. DFM microscopy of *in vitro* cultivated *TPA* Nichols incubated for seven days with 10% rabbit α-Tpp17.**

**S4 Movie. DFM microscopy of *in vitro* cultivated *TPA* Nichols incubated for seven days with 10% rabbit α-TP0751.**

**S5 Movie. DFM microscopy of *in vitro* cultivated *TPA* Nichols incubated for seven days with 10% rabbit α-*Pf*Trx^BamA/ECL4^.**

**S6 Movie. DFM microscopy of *in vitro* cultivated *TPA* Nichols incubated for seven days with 5% pooled MSS.**

**S7 Movie. DFM microscopy of *in vitro* cultivated *TPA* incubated for seven days with 5% pooled NMS.**

**S8 Movie. DFM microscopy of *in vitro* cultivated *TPA* Nichols incubated for seven days with 5% pooled mouse α-Tpp17.**

**S9 Movie. DFM microscopy of *in vitro* cultivated *TPA* Nichols incubated for seven days with 5% pooled mouse α-TP0751.**

**S10 Movie. DFM microscopy of *in vitro* cultivated *TPA* Nichols incubated for seven days with 5% pooled mouse α-*Pf*Trx^BamA/ECL4^.**

## References

1. Ghanem KG, Ram S, Rice PA. The modern epidemic of syphilis. N Engl J Med. 2020;382(9):845–54. doi: 10.1056/NEJMra1901593. PubMed PMID: 32101666.

2. Peeling RW, Mabey D, Chen XS, Garcia PJ. Syphilis. Lancet. 2023;402(10398):336–46. doi: 10.1016/s0140-6736(22)02348-0. PubMed PMID: 37481272.

3. Kojima N, Klausner JD. An update on the global epidemiology of syphilis. Curr Epidemiol Rep. 2018;5(1):24–38. Epub 2018/08/18. doi: 10.1007/s40471-018-0138-z. PubMed PMID: 30116697; PubMed Central PMCID: PMCPMC6089383.

4. Gottlieb SL, Deal CD, Giersing B, Rees H, Bolan G, Johnston C, et al. The global roadmap for advancing development of vaccines against sexually transmitted infections: Update and next steps. Vaccine. 2016;34(26):2939–47. doi: 10.1016/j.vaccine.2016.03.111. PubMed PMID: 27105564.

5. Kojima N, Konda KA, Klausner JD. Notes on syphilis vaccine development. Front Immunol. 2022;13:952284. Epub 20220728. doi: 10.3389/fimmu.2022.952284. PubMed PMID: 35967432; PubMed Central PMCID: PMCPMC9365935.

6. Sell S, Norris SJ. The biology, pathology, and immunology of syphilis. Int Rev Exp Pathol. 1983;24:203–76. PubMed PMID: 6840996.

7. Schell RF. Rabbit and hamster models of treponemal infection. In: Schell RF, Musher DM, editors. Pathogenesis and immunology of Treponemal infection. New York: Marcel Dekker, Inc.; 1983. p. 121–35.

8. Turner TB, Hollander DH. Biology of the Treponematoses. Geneva World Health Organziation; 1957.

9. Magnuson HJ, Rosenau BJ. The rate of development and degree of acquired immunity in experimental syphilis. Am J Syph Gonorrhea Vener Dis. 1948;32(5):418–36. Epub 1948/09/01. PubMed PMID: 18876783.

10. Lukehart SA. Scientific monogamy: thirty years dancing with the same bug: 2007 Thomas Parran Award Lecture. Sex Transm Dis. 2008;35(1):2–7. doi: 10.1097/OLQ.0b013e318162c4f2. PubMed PMID: 18157060.

11. Marra CM, Tantalo LC, Sahi SK, Dunaway SB, Lukehart SA. Reduced *Treponema pallidum*-specific opsonic antibody activity in HIV-infected patients with syphilis. The Journal of Infectious Diseases. 2015;213(8):1348–54. doi: 10.1093/infdis/jiv591.

12. Cruz AR, Ramirez LG, Zuluaga AV, Pillay A, Abreu C, Valencia CA, et al. Immune evasion and recognition of the syphilis spirochete in blood and skin of secondary syphilis patients: two immunologically distinct compartments. PLoS Negl Trop Dis. 2012;6(7):e1717. Epub 20120717. doi: 10.1371/journal.pntd.0001717. PubMed PMID: 22816000; PubMed Central PMCID: PMCPMC3398964.

13. Hawley KL, Cruz AR, Benjamin SJ, La Vake CJ, Cervantes JL, LeDoyt M, et al. IFNγ enhances CD64-potentiated phagocytosis of *Treponema pallidum* opsonized with human syphilitic serum by human macrophages. Front Immunol. 2017;8:1227. Epub 20171005. doi: 10.3389/fimmu.2017.01227. PubMed PMID: 29051759; PubMed Central PMCID: PMCPMC5633599.

14. Radolf JD, Lukehart SA. Immunology of Syphilis. In: Radolf JD, Lukehart SA, editors. Pathogenic Treponemes: Cellular and molecular biology. Norfolk, UK: Caister Academic Press; 2006. p. 285–322.

15. Folds JD, Rauchbach AS, Shores E, Saunders JM. Evaluation of the inbred mouse as a model for experimental *Treponema pallidum* infection. Scand J Immunol. 1983;18(3):201–6. doi: 10.1111/j.1365-3083.1983.tb00858.x. PubMed PMID: 6353562.

16. Gueft B, Rosahn PD. Experimental mouse syphilis, a critical review of the literature. Am J Syph Gonorrhea Vener Dis. 1948;32(1):59–88. PubMed PMID: 18917627.

17. Silver AC, Dunne DW, Zeiss CJ, Bockenstedt LK, Radolf JD, Salazar JC, et al. MyD88 deficiency markedly worsens tissue inflammation and bacterial clearance in mice infected with *Treponema pallidum*, the agent of syphilis. PLoS One. 2013;8(8):e71388. doi: 10.1371/journal.pone.0071388. PubMed PMID: 23940747; PubMed Central PMCID: PMCPMC3734110.

18. Ferguson MR, Delgado KN, McBride S, Orbe IC, La Vake CJ, Caimano MJ, et al. Use of Epivolve phage display to generate a monoclonal antibody with opsonic activity directed against a subdominant epitope on extracellular loop 4 of *Treponema pallidum* BamA (TP0326). Frontiers in Immunology. 2023;14. doi: 10.3389/fimmu.2023.1222267.

19. Rosahn PD, Gueft B, Rowe CL. Experimental mouse syphilis; organ distribution of the infectious agent. Am J Syph Gonorrhea Vener Dis. 1948;32(4):327–36. PubMed PMID: 18867296.

20. Cox DL, Luthra A, Dunham-Ems S, Desrosiers DC, Salazar JC, Caimano MJ, et al. Surface immunolabeling and consensus computational framework to identify candidate rare outer membrane proteins of *Treponema pallidum*. Infect Immun. 2010;78(12):5178–94. Epub 20100927. doi: 10.1128/iai.00834-10. PubMed PMID: 20876295; PubMed Central PMCID: PMCPMC2981305.

21. Liu J, Howell JK, Bradley SD, Zheng Y, Zhou ZH, Norris SJ. Cellular architecture of *Treponema pallidum*: novel flagellum, periplasmic cone, and cell envelope as revealed by cryo electron tomography. J Mol Biol. 2010;403(4):546–61. doi: 10.1016/j.jmb.2010.09.020. PubMed PMID: 20850455; PubMed Central PMCID: PMCPMC2957517.

22. Radolf JD, Deka RK, Anand A, Smajs D, Norgard MV, Yang XF. *Treponema pallidum*, the syphilis spirochete: making a living as a stealth pathogen. Nat Rev Microbiol. 2016. doi: 10.1038/nrmicro.2016.141. PubMed PMID: 27721440.

23. Radolf JD, Kumar S. The *Treponema pallidum* outer membrane. Curr Top Microbiol Immunol. 2018;415:1–38. Epub 2017/08/30. doi: 10.1007/82_2017_44. PubMed PMID: 28849315; PubMed Central PMCID: PMCPMC5924592.

24. Hawley KL, Montezuma-Rusca JM, Delgado KN, Singh N, Uversky VN, Caimano MJ, et al. Structural modeling of the *Treponema pallidum* OMPeome: a roadmap for deconvolution of syphilis pathogenesis and development of a syphilis vaccine. J Bacteriol. 2021;203(15):e0008221. Epub 2021/05/12. doi: 10.1128/JB.00082-21. PubMed PMID: 33972353; PubMed Central PMCID: PMCPMC8407342.

25. Ávila-Nieto C, Pedreño-López N, Mitjà O, Clotet B, Blanco J, Carrillo J. Syphilis vaccine: challenges, controversies and opportunities. Front Immunol. 2023;14:1126170. Epub 20230406. doi: 10.3389/fimmu.2023.1126170. PubMed PMID: 37090699; PubMed Central PMCID: PMCPMC10118025.

26. von Kügelgen A, van Dorst S, Alva V, Bharat TAM. A multidomain connector links the outer membrane and cell wall in phylogenetically deep-branching bacteria. Proc Natl Acad Sci U S A. 2022;119(33):e2203156119. Epub 20220809. doi: 10.1073/pnas.2203156119. PubMed PMID: 35943982; PubMed Central PMCID: PMCPMC9388160.

27. Delgado KN, Montezuma-Rusca JM, Orbe IC, Caimano MJ, La Vake CJ, Luthra A, et al. Extracellular loops of the *Treponema pallidum* FadL orthologs TP0856 and TP0858 elicit IgG antibodies and IgG(+)-specific B-cells in the rabbit model of experimental syphilis. mBio. 2022;13(4):e0163922. Epub 20220712. doi: 10.1128/mbio.01639-22. PubMed PMID: 35862766; PubMed Central PMCID: PMCPMC9426418.

28. Cappelli L, Cinelli P, Perrotta A, Veggi D, Audagnotto M, Tuscano G, et al. Computational structure-based approach to study chimeric antigens using a new protein scaffold displaying foreign epitopes. Faseb j. 2024;38(1):e23326. doi: 10.1096/fj.202202130R. PubMed PMID: 38019196.

29. Fegan JE, Calmettes C, Islam EA, Ahn SK, Chaudhuri S, Yu RH, et al. Utility of hybrid transferrin binding protein antigens for protection against pathogenic *Neisseria* species. Front Immunol. 2019;10:247. Epub 20190219. doi: 10.3389/fimmu.2019.00247. PubMed PMID: 30837995; PubMed Central PMCID: PMCPMC6389628.

30. Collar AL, Linville AC, Core SB, Wheeler CM, Geisler WM, Peabody DS, et al. Antibodies to variable domain 4 linear epitopes of the *Chlamydia trachomatis* major outer membrane protein are not associated with *Chlamydia* resolution or reinfection in women. mSphere. 2020;5(5). Epub 2020/09/25. doi: 10.1128/mSphere.00654-20. PubMed PMID: 32968007; PubMed Central PMCID: PMCPMC7568647.

31. Spagnoli G, Pouyanfard S, Cavazzini D, Canali E, Maggi S, Tommasino M, et al. Broadly neutralizing antiviral responses induced by a single-molecule HPV vaccine based on thermostable thioredoxin-L2 multiepitope nanoparticles. Sci Rep. 2017;7(1):18000. Epub 2017/12/23. doi: 10.1038/s41598-017-18177-1. PubMed PMID: 29269879; PubMed Central PMCID: PMCPMC5740060.

32. Edmondson DG, Hu B, Norris SJ. Long-term *in vitro* culture of the syphilis spirochete *Treponema pallidum* subsp. *pallidum*. mBio. 2018;9(3). Epub 2018/06/28. doi: 10.1128/mBio.01153-18. PubMed PMID: 29946052; PubMed Central PMCID: PMCPMC6020297.

33. Du Z, Su H, Wang W, Ye L, Wei H, Peng Z, et al. The trRosetta server for fast and accurate protein structure prediction. Nat Protoc. 2021;16(12):5634–51. Epub 20211110. doi: 10.1038/s41596-021-00628-9. PubMed PMID: 34759384.

34. Abramson J, Adler J, Dunger J, Evans R, Green T, Pritzel A, et al. Accurate structure prediction of biomolecular interactions with AlphaFold 3. Nature. 2024. Epub 20240508. doi: 10.1038/s41586-024-07487-w. PubMed PMID: 38718835.

35. van den Berg B. Bacterial cleanup: lateral diffusion of hydrophobic molecules through protein channel walls. Biomol Concepts. 2010;1(3-4):263–70. doi: 10.1515/bmc.2010.024. PubMed PMID: 25962002.

36. van den Berg B, Black PN, Clemons WM, Jr., Rapoport TA. Crystal structure of the long-chain fatty acid transporter FadL. Science. 2004;304(5676):1506-9. doi: 10.1126/science.1097524. PubMed PMID: 15178802.

37. Koronakis V, Sharff A, Koronakis E, Luisi B, Hughes C. Crystal structure of the bacterial membrane protein TolC central to multidrug efflux and protein export. Nature. 2000;405(6789):914-9. doi: 10.1038/35016007. PubMed PMID: 10879525.

38. Lei HT, Chou TH, Su CC, Bolla JR, Kumar N, Radhakrishnan A, et al. Crystal structure of the open state of the *Neisseria gonorrhoeae* MtrE outer membrane channel. PLoS One. 2014;9(6):e97475. Epub 20140605. doi: 10.1371/journal.pone.0097475. PubMed PMID: 24901251; PubMed Central PMCID: PMCPMC4046963.

39. Ponomarenko J, Bui H-H, Li W, Fusseder N, Bourne PE, Sette A, et al. ElliPro: a new structure-based tool for the prediction of antibody epitopes. BMC Bioinformatics. 2008;9(1):514. doi: 10.1186/1471-2105-9-514.

40. Kringelum JV, Lundegaard C, Lund O, Nielsen M. Reliable B cell epitope predictions: impacts of method development and improved benchmarking. PLoS Comput Biol. 2012;8(12):e1002829. Epub 20121227. doi: 10.1371/journal.pcbi.1002829. PubMed PMID: 23300419; PubMed Central PMCID: PMCPMC3531324.

41. Hanff PA, Bishop NH, Miller JN, Lovett MA. Humoral immune response in experimental syphilis to polypeptides of *Treponema pallidum*. J Immunol. 1983;131(4):1973–7. PubMed PMID: 6352809.

42. Chamberlain NR, Brandt ME, Erwin AL, Radolf JD, Norgard MV. Major integral membrane protein immunogens of *Treponema pallidum* are proteolipids. Infection and Immunity. 1989;57(9):2872–7. doi: 10.1128/iai.57.9.2872-2877.1989.

43. Lieberman NAP, Lin MJ, Xie H, Shrestha L, Nguyen T, Huang ML, et al. *Treponema pallidum* genome sequencing from six continents reveals variability in vaccine candidate genes and dominance of Nichols clade strains in Madagascar. PLoS Negl Trop Dis. 2021;15(12):e0010063. Epub 20211222. doi: 10.1371/journal.pntd.0010063. PubMed PMID: 34936652; PubMed Central PMCID: PMCPMC8735616.

44. Arora N, Schuenemann VJ, Jäger G, Peltzer A, Seitz A, Herbig A, et al. Origin of modern syphilis and emergence of a pandemic *Treponema pallidum* cluster. Nat Microbiol. 2016;2:16245. Epub 20161205. doi: 10.1038/nmicrobiol.2016.245. PubMed PMID: 27918528.

45. Beale MA, Marks M, Cole MJ, Lee MK, Pitt R, Ruis C, et al. Global phylogeny of *Treponema pallidum* lineages reveals recent expansion and spread of contemporary syphilis. Nat Microbiol. 2021;6(12):1549–60. Epub 20211124. doi: 10.1038/s41564-021-01000-z. PubMed PMID: 34819643; PubMed Central PMCID: PMCPMC8612932.

46. Luthra A, Anand A, Hawley KL, LeDoyt M, La Vake CJ, Caimano MJ, et al. A homology model reveals novel structural features and an immunodominant surface loop/opsonic target in the *Treponema pallidum* BamA ortholog TP_0326. J Bacteriol. 2015;197(11):1906–20. doi: 10.1128/JB.00086-15. PubMed PMID: 25825429; PubMed Central PMCID: PMCPMC4420902.

47. Luthra A, Montezuma-Rusca JM, La Vake CJ, LeDoyt M, Delgado KN, Davenport TC, et al. Evidence that immunization with TP0751, a bipartite *Treponema pallidum* lipoprotein with an intrinsically disordered region and lipocalin fold, fails to protect in the rabbit model of experimental syphilis. PLoS Pathog. 2020;16(9):e1008871. Epub 20200916. doi: 10.1371/journal.ppat.1008871. PubMed PMID: 32936831; PubMed Central PMCID: PMCPMC7521688.

48. Edmondson DG, Norris SJ. *In vitro* cultivation of the syphilis spirochete *Treponema pallidum*. Curr Protoc. 2021;1(2):e44. doi: 10.1002/cpz1.44. PubMed PMID: 33599121; PubMed Central PMCID: PMCPMC7986111.

49. Vij R, Lin Z, Chiang N, Vernes JM, Storek KM, Park S, et al. A targeted boost-and-sort immunization strategy using *Escherichia coli* BamA identifies rare growth inhibitory antibodies. Sci Rep. 2018;8(1):7136. Epub 20180508. doi: 10.1038/s41598-018-25609-z. PubMed PMID: 29740124; PubMed Central PMCID: PMCPMC5940829.

50. De Lay BD, Cameron TA, De Lay NR, Norris SJ, Edmondson DG. Comparison of transcriptional profiles of *Treponema pallidum* during experimental infection of rabbits and *in vitro* culture: Highly similar, yet different. PLoS Pathog. 2021;17(9):e1009949. Epub 20210927. doi: 10.1371/journal.ppat.1009949. PubMed PMID: 34570834; PubMed Central PMCID: PMCPMC8525777.

51. Houston S, Gomez A, Geppert A, Goodyear MC, Cameron CE. In-Depth proteome coverage of *in vitro*-cultured *Treponema pallidum* and quantitative comparison analyses with *in vivo*-grown treponemes. J Proteome Res. 2024;23(5):1725–43. Epub 20240418. doi: 10.1021/acs.jproteome.3c00891. PubMed PMID: 38636938.

52. Houston S, Gomez A, Geppert A, Eshghi A, Smith DS, Waugh S, et al. Deep proteome coverage advances knowledge of *Treponema pallidum* protein expression profiles during infection. Sci Rep. 2023;13(1):18259. Epub 20231025. doi: 10.1038/s41598-023-45219-8. PubMed PMID: 37880309; PubMed Central PMCID: PMCPMC10600179.

53. McGill MA, Edmondson DG, Carroll JA, Cook RG, Orkiszewski RS, Norris SJ. Characterization and serologic analysis of the *Treponema pallidum* proteome. Infect Immun. 2010;78(6):2631–43. doi: 10.1128/IAI.00173-10. PubMed PMID: 20385758; PubMed Central PMCID: PMCPMC2876534.

54. Osbak KK, Houston S, Lithgow KV, Meehan CJ, Strouhal M, Smajs D, et al. Characterizing the syphilis-causing *Treponema pallidum* ssp. *pallidum* proteome using complementary mass spectrometry. PLoS Negl Trop Dis. 2016;10(9):e0004988. doi: 10.1371/journal.pntd.0004988. PubMed PMID: 27606673; PubMed Central PMCID: PMCPMC5015957.

55. Zepp F. Principles of vaccine design-Lessons from nature. Vaccine. 2010;28 Suppl 3:C14–24. Epub 2010/08/18. doi: 10.1016/j.vaccine.2010.07.020. PubMed PMID: 20713252.

56. Sanchez-Trincado JL, Gomez-Perosanz M, Reche PA. Fundamentals and methods for T- and B-cell epitope prediction. Journal of Immunology Research. 2017;2017:2680160. doi: 10.1155/2017/2680160.

57. Matthias KA, Strader MB, Nawar HF, Gao YS, Lee J, Patel DS, et al. Heterogeneity in non-epitope loop sequence and outer membrane protein complexes alters antibody binding to the major porin protein PorB in serogroup B *Neisseria meningitidis*. Mol Microbiol. 2017;105(6):934–53. doi: 10.1111/mmi.13747. PubMed PMID: 28708335.

58. Noinaj N, Kuszak AJ, Balusek C, Gumbart JC, Buchanan SK. Lateral opening and exit pore formation are required for BamA function. Structure. 2014;22(7):1055–62. Epub 20140626. doi: 10.1016/j.str.2014.05.008. PubMed PMID: 24980798; PubMed Central PMCID: PMCPMC4100585.

59. Pautsch A, Schulz GE. High-resolution structure of the OmpA membrane domain. J Mol Biol. 2000;298(2):273–82. doi: 10.1006/jmbi.2000.3671. PubMed PMID: 10764596.

60. Straatsma TP, Soares TA. Characterization of the outer membrane protein OprF of *Pseudomonas aeruginosa* in a lipopolysaccharide membrane by computer simulation. Proteins. 2009;74(2):475–88. doi: 10.1002/prot.22165. PubMed PMID: 18655068; PubMed Central PMCID: PMCPMC2610247.

61. Horst R, Stanczak P, Wuthrich K. NMR polypeptide backbone conformation of the *E. coli* outer membrane protein W. Structure. 2014;22(8):1204–9. doi: 10.1016/j.str.2014.05.016. PubMed PMID: 25017731; PubMed Central PMCID: PMCPMC4150354.

62. Oomen CJ, Hoogerhout P, Kuipers B, Vidarsson G, van Alphen L, Gros P. Crystal structure of an anti-meningococcal subtype P1.4 PorA antibody provides basis for peptide-vaccine design. J Mol Biol. 2005;351(5):1070–80. Epub 2005/07/26. doi: 10.1016/j.jmb.2005.06.061. PubMed PMID: 16038932.

63. Domínguez-Medina CC, Pérez-Toledo M, Schager AE, Marshall JL, Cook CN, Bobat S, et al. Outer membrane protein size and LPS O-antigen define protective antibody targeting to the *Salmonella* surface. Nat Commun. 2020;11(1):851. Epub 20200212. doi: 10.1038/s41467-020-14655-9. PubMed PMID: 32051408; PubMed Central PMCID: PMCPMC7015928.

64. Tifrea DF, Pal S, Fairman J, Massari P, de la Maza LM. Protection against a chlamydial respiratory challenge by a chimeric vaccine formulated with the *Chlamydia muridarum* major outer membrane protein variable domains using the *Neisseria* lactamica porin B as a scaffold. NPJ Vaccines. 2020;5:37. Epub 2020/05/16. doi: 10.1038/s41541-020-0182-9. PubMed PMID: 32411400; PubMed Central PMCID: PMCPMC7210953.

65. Cia G, Pucci F, Rooman M. Critical review of conformational B-cell epitope prediction methods. Brief Bioinform. 2023;24(1). doi: 10.1093/bib/bbac567. PubMed PMID: 36611255.

66. Radolf JD, Robinson EJ, Bourell KW, Akins DR, Porcella SF, Weigel LM, et al. Characterization of outer membranes isolated from *Treponema pallidum*, the syphilis spirochete. Infect Immun. 1995;63(11):4244–52. PubMed PMID: 7591054; PubMed Central PMCID: PMCPMC173603.

67. Radolf JD, Norgard MV, Schulz WW. Outer membrane ultrastructure explains the limited antigenicity of virulent *Treponema pallidum*. Proc Natl Acad Sci U S A. 1989;86(6):2051–5. PubMed PMID: 2648388; PubMed Central PMCID: PMCPMC286845.

68. Cox DL, Chang P, McDowall AW, Radolf JD. The outer membrane, not a coat of host proteins, limits antigenicity of virulent *Treponema pallidum*. Infect Immun. 1992;60(3):1076–83. doi: 10.1128/iai.60.3.1076-1083.1992. PubMed PMID: 1541522; PubMed Central PMCID: PMCPMC257596.

69. Plotkin SA, Plotkin SA. Correlates of Vaccine-Induced Immunity. Clinical Infectious Diseases. 2008;47(3):401–9. doi: 10.1086/589862.

70. Alder JD, Friess L, Tengowski M, Schell RF. Phagocytosis of opsonized T*reponema pallidum subsp. pallidum* proceeds slowly. Infect Immun. 1990;58(5):1167–73. doi: 10.1128/iai.58.5.1167-1173.1990. PubMed PMID: 2182536; PubMed Central PMCID: PMCPMC258605.

71. Lukehart SA, Shaffer JM, Baker-Zander SA. A subpopulation of *Treponema pallidum* is resistant to phagocytosis: possible mechanism of persistence. J Infect Dis. 1992;166(6):1449–53. doi: 10.1093/infdis/166.6.1449. PubMed PMID: 1431264.

72. Bourell KW, Schulz W, Norgard MV, Radolf JD. *Treponema pallidum* rare outer membrane proteins: analysis of mobility by freeze-fracture electron microscopy. J Bacteriol. 1994;176(6):1598–608. doi: 10.1128/jb.176.6.1598-1608.1994. PubMed PMID: 8132453; PubMed Central PMCID: PMCPMC205244.

73. Duchemin AM, Ernst LK, Anderson CL. Clustering of the high affinity Fc receptor for immunoglobulin G (Fc gamma RI) results in phosphorylation of its associated gamma-chain. J Biol Chem. 1994;269(16):12111–7. PubMed PMID: 7512959.

74. Nelson RA, Jr., Mayer MM. Immobilization of *Treponema pallidum in vitro* by antibody produced in syphilitic infection. J Exp Med. 1949;89(4):369–93. Epub 1949/04/01. PubMed PMID: 18113911; PubMed Central PMCID: PMC2135874.

75. Bishop NH, Miller JN. Humoral immunity in experimental syphilis. I. The demonstration of resistance conferred by passive immunization. J Immunol. 1976;117(1):191–6. PubMed PMID: 778261.

76. Azadegan AA, Tabor DR, Schell RF, LeFrock JL. Cobra venom factor abrogates passive humoral resistance to syphilitic infection in hamsters. Infect Immun. 1984;44(3):740–2. doi: 10.1128/iai.44.3.740-742.1984. PubMed PMID: 6724696; PubMed Central PMCID: PMCPMC263686.

77. Weiser RS, Erickson D, Perine PL, Pearsall NN. Immunity to syphilis: passive transfer in rabbits using serial doses of immune serum. Infect Immun. 1976;13(5):1402–7. Epub 1976/05/01. doi: 10.1128/iai.13.5.1402-1407.1976. PubMed PMID: 773833; PubMed Central PMCID: PMCPMC420773.

78. Perine PL, Weiser RS, Klebanoff SJ. Immunity to syphilis. I. Passive transfer in rabbits with hyperimmune serum. Infect Immun. 1973;8(5):787–90. PubMed PMID: 4584052; PubMed Central PMCID: PMCPMC422928.

79. Fitzgerald TJ, Miller JN, Sykes JA. *Treponema pallidum* (Nichols strain) in tissue cultures: cellular attachment, entry, and survival. Infect Immun. 1975;11(5):1133–40. PubMed PMID: 1091562; PubMed Central PMCID: PMCPMC415188.

80. Fitzgerald TJ, Repesh LA, Blanco DR, Miller JN. Attachment of *Treponema pallidum* to fibronectin, laminin, collagen IV, and collagen I, and blockage of attachment by immune rabbit IgG. Br J Vener Dis. 1984;60(6):357–63. PubMed PMID: 6394096; PubMed Central PMCID: PMCPMC1046381.

81. Izard J, Renken C, Hsieh CE, Desrosiers DC, Dunham-Ems S, La Vake C, et al. Cryo-electron tomography elucidates the molecular architecture of *Treponema pallidum*, the syphilis spirochete. J Bacteriol. 2009;191(24):7566–80. doi: 10.1128/JB.01031-09. PubMed PMID: 19820083; PubMed Central PMCID: PMCPMC2786590.

82. Cameron CE, Brouwer NL, Tisch LM, Kuroiwa JM. Defining the interaction of the *Treponema pallidum* adhesin Tp0751 with laminin. Infect Immun. 2005;73(11):7485–94. doi: 10.1128/iai.73.11.7485-7494.2005. PubMed PMID: 16239550; PubMed Central PMCID: PMCPMC1273862.

83. Houston S, Hof R, Francescutti T, Hawkes A, Boulanger MJ, Cameron CE. Bifunctional role of the *Treponema pallidum* extracellular matrix binding adhesin Tp0751. Infect Immun. 2011;79(3):1386–98. Epub 20101213. doi: 10.1128/iai.01083-10. PubMed PMID: 21149586; PubMed Central PMCID: PMCPMC3067502.

84. Haynes AM, Godornes C, Ke W, Giacani L. Evaluation of the protective ability of the *Treponema pallidum subsp. pallidum* Tp0126 OmpW homolog in the rabbit model of syphilis. Infect Immun. 2019;87(8). Epub 20190723. doi: 10.1128/iai.00323-19. PubMed PMID: 31182617; PubMed Central PMCID: PMCPMC6652746.

85. Centurion-Lara A, Castro C, Barrett L, Cameron C, Mostowfi M, Van Voorhis WC, et al. *Treponema pallidum* major sheath protein homologue TprK is a target of opsonic antibody and the protective immune response. J Exp Med. 1999;189(4):647–56. PubMed PMID: 9989979; PubMed Central PMCID: PMCPMC2192927.

86. Cameron CE, Lukehart SA, Castro C, Molini B, Godornes C, Van Voorhis WC. Opsonic potential, protective capacity, and sequence conservation of the *Treponema pallidum* subspecies *pallidum* Tp92. J Infect Dis. 2000;181(4):1401–13. doi: 10.1086/315399. PubMed PMID: 10762571.

87. Sun ES, Molini BJ, Barrett LK, Centurion-Lara A, Lukehart SA, Van Voorhis WC. Subfamily I *Treponema pallidum* repeat protein family: sequence variation and immunity. Microbes Infect. 2004;6(8):725–37. Epub 2004/06/23. doi: 10.1016/j.micinf.2004.04.001. PubMed PMID: 15207819.

88. Noinaj N, Gumbart JC, Buchanan SK. The β-barrel assembly machinery in motion. Nat Rev Microbiol. 2017;15(4):197–204. doi: 10.1038/nrmicro.2016.191. PubMed PMID: 28216659; PubMed Central PMCID: PMCPMC5455337.

89. Oleastro M, Menard A. The role of *Helicobacter pylori* outer membrane proteins in adherence and pathogenesis. Biology (Basel). 2013;2(3):1110–34. Epub 2013/01/01. doi: 10.3390/biology2031110. PubMed PMID: 24833057; PubMed Central PMCID: PMCPMC3960876.

90. Krishnan S, Prasadarao NV. Outer membrane protein A and OprF: versatile roles in Gram-negative bacterial infections. FEBS J. 2012;279(6):919–31. Epub 2012/01/14. doi: 10.1111/j.1742-4658.2012.08482.x. PubMed PMID: 22240162; PubMed Central PMCID: PMCPMC3338869.

91. Azghani AO, Idell S, Bains M, Hancock RE. *Pseudomonas aeruginosa* outer membrane protein F is an adhesin in bacterial binding to lung epithelial cells in culture. Microb Pathog. 2002;33(3):109–14. doi: 10.1006/mpat.2002.0514. PubMed PMID: 12220987.

92. Fairman JW, Dautin N, Wojtowicz D, Liu W, Noinaj N, Barnard TJ, et al. Crystal structures of the outer membrane domain of intimin and invasin from enterohemorrhagic *E. coli* and enteropathogenic *Y. pseudotuberculosis*. Structure. 2012;20(7):1233–43. Epub 20120531. doi: 10.1016/j.str.2012.04.011. PubMed PMID: 22658748; PubMed Central PMCID: PMCPMC3392549.

93. Sievers F, Wilm A, Dineen D, Gibson TJ, Karplus K, Li W, et al. Fast, scalable generation of high-quality protein multiple sequence alignments using Clustal Omega. Mol Syst Biol. 2011;7:539. Epub 2011/10/13. doi: 10.1038/msb.2011.75. PubMed PMID: 21988835; PubMed Central PMCID: PMCPMC3261699.

94. Joshi NA FJ. Sickle: A sliding-window, adaptive, quality-based trimming tool for FastQ files (Version 1.33) 2011. Available from: https://github.com/najoshi/sickle.

95. Magoc T, Wood D, Salzberg SL. EDGE-pro: Estimated Degree of Gene Expression in Prokaryotic Genomes. Evol Bioinform Online. 2013;9:127–36. Epub 20130310. doi: 10.4137/ebo.S11250. PubMed PMID: 23531787; PubMed Central PMCID: PMCPMC3603529.

96. Wagner GP, Kin K, Lynch VJ. Measurement of mRNA abundance using RNA-seq data: RPKM measure is inconsistent among samples. Theory Biosci. 2012;131(4):281–5. Epub 20120808. doi: 10.1007/s12064-012-0162-3. PubMed PMID: 22872506.

