## Supplemental Figures for "Immunodominant extracellular loops of *Treponema pallidum* FadL outer membrane proteins elicit antibodies with opsonic and growth-inhibitory activities"

### Supporting Information:

**S1A** and **S1B Fig.**

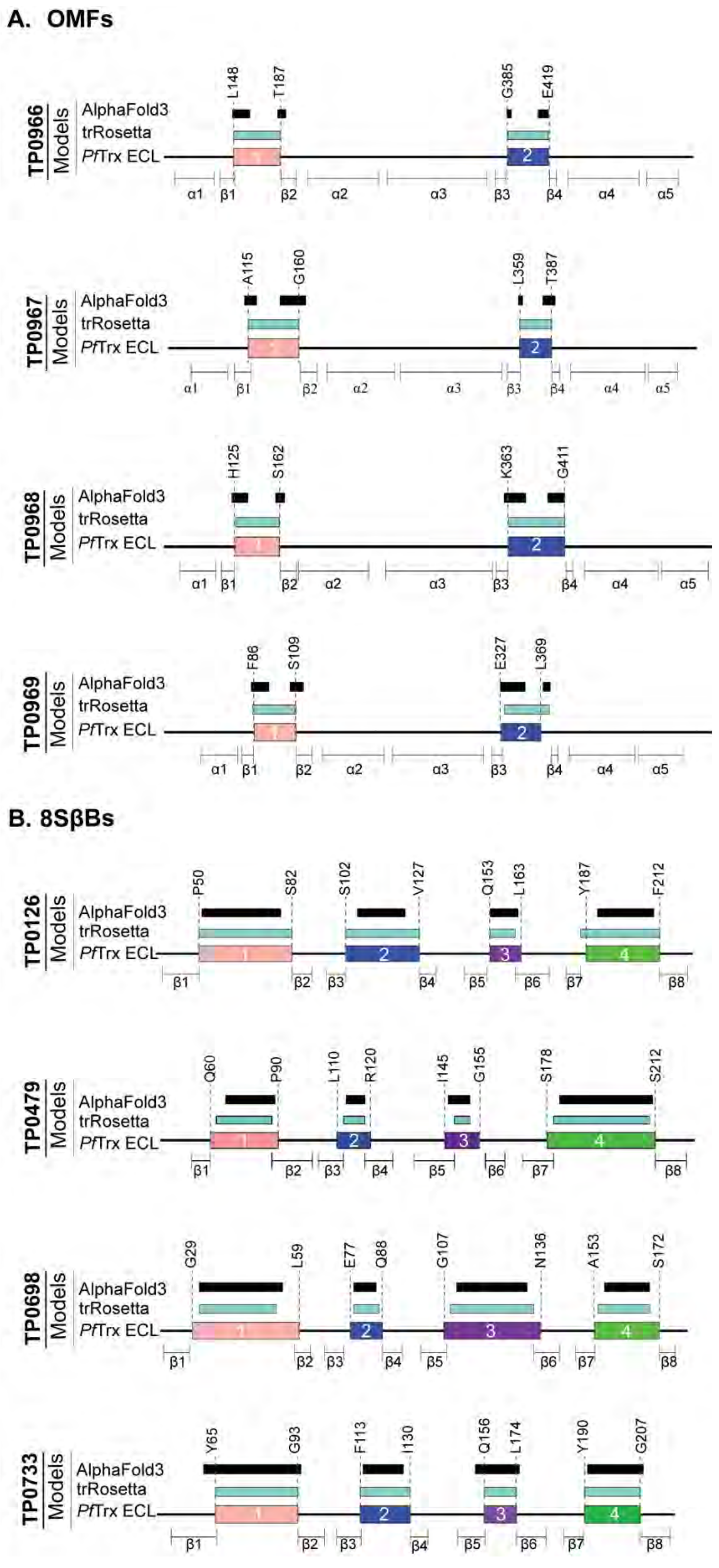

**S1C Fig.**

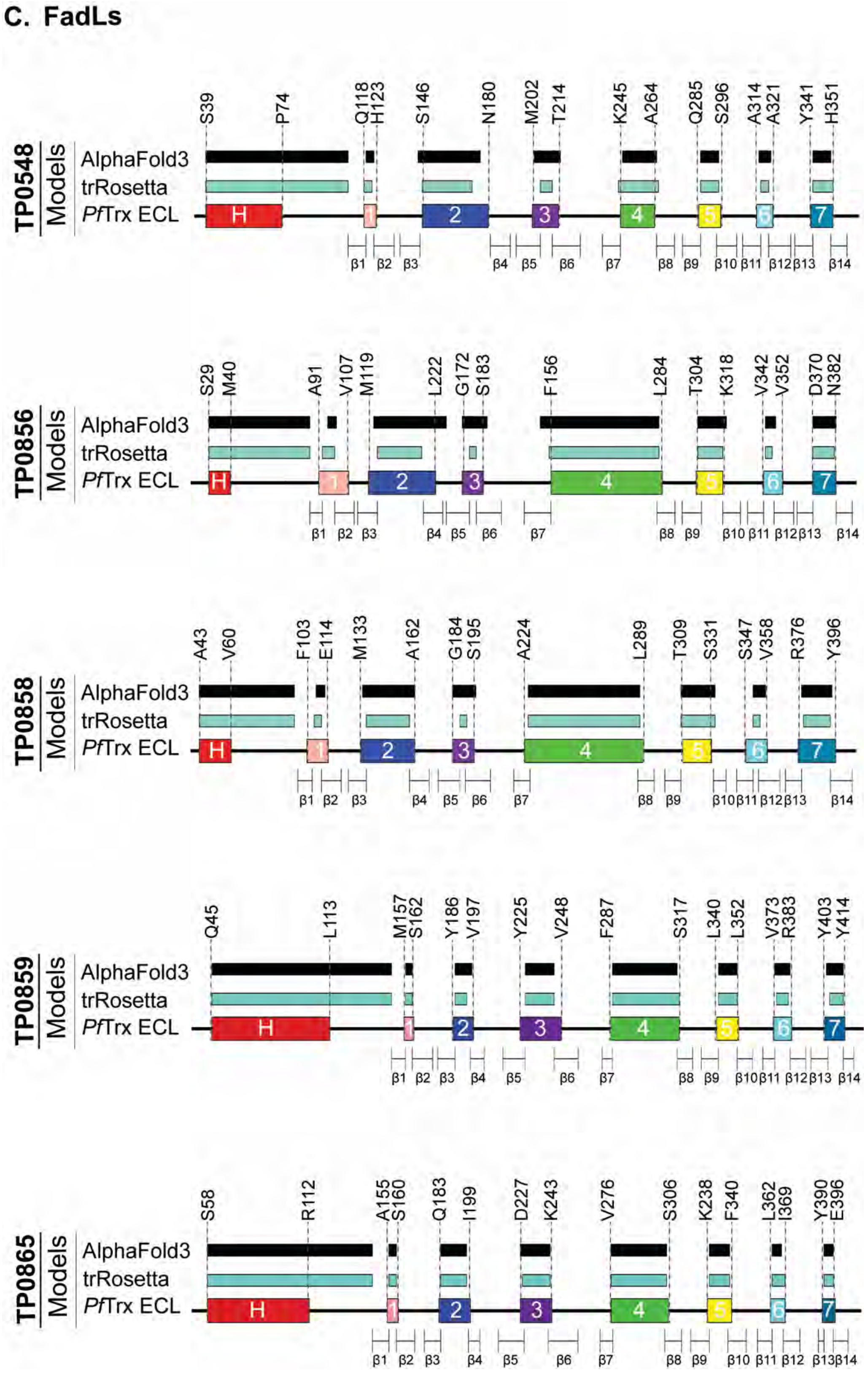
C. FadLs
TP0548
Models
AlphaFold3
trRosetta
PfTrx ECL
S39 P74 Q118 H123 S146 N180 M202 T214 K245 A264 Q285 S296 A314 A321 Y341 H351
H 1 2 3 4 5 6 7
β1 β2 β3 β4 β5 β6 β7 β8 β9 β10 β11 β12 β13 β14
TP0856
Models
AlphaFold3
trRosetta
PfTrx ECL
S29 M40 A91 V107 M119 L222 G172 S183 F156 L284 T304 K318 V342 V352 D370 N382
H 1 2 3 4 5 6 7
β1 β2 β3 β4 β5 β6 β7 β8 β9 β10 β11 β12 β13 β14
TP0858
Models
AlphaFold3
trRosetta
PfTrx ECL
A43 V60 F103 E114 M133 A162 G184 S195 A224 L289 T309 S331 S347 V358 R376 Y396
H 1 2 3 4 5 6 7
β1 β2 β3 β4 β5 β6 β7 β8 β9 β10 β11 β12 β13 β14
TP0859
Models
AlphaFold3
trRosetta
PfTrx ECL
Q45 L113 M157 S162 Y186 V197 Y225 V248 F287 S317 L340 L352 V373 R383 Y403 Y414
H 1 2 3 4 5 6 7
β1 β2 β3 β4 β5 β6 β7 β8 β9 β10 β11 β12 β13 β14
TP0865
Models
AlphaFold3
trRosetta
PfTrx ECL
S58 R112 A155 S160 Q183 I199 D227 K243 V276 S306 K238 F340 L362 I369 Y390 E396
H 1 2 3 4 5 6 7
β1 β2 β3 β4 β5 β6 β7 β8 β9 β10 β11 β12 β13 β14

**S2 Fig.**

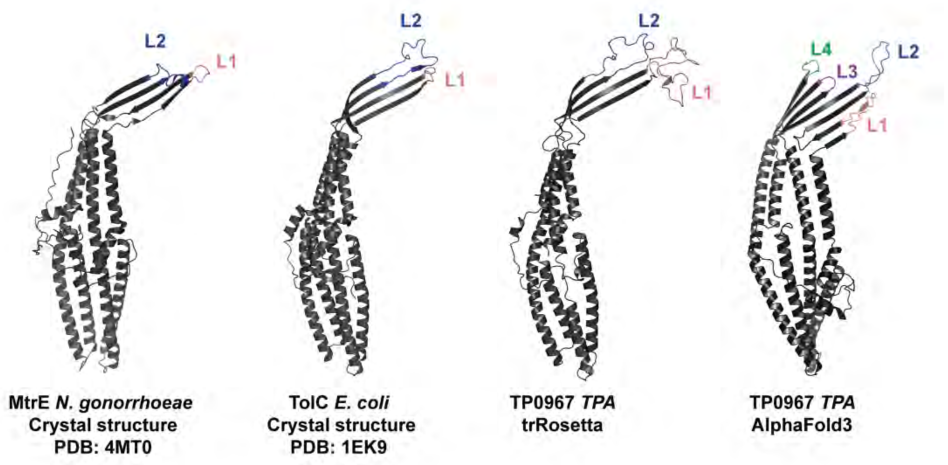
L2
L1
L2
L1
L2
L1
L4
L3
L2
L1
MtrE *N. gonorrhoeae*
Crystal structure
PDB: 4MT0
TolC *E. coli*
Crystal structure
PDB: 1EK9
TP0967 *TPA*
trRosetta
TP0967 *TPA*
AlphaFold3

### S3 Fig.

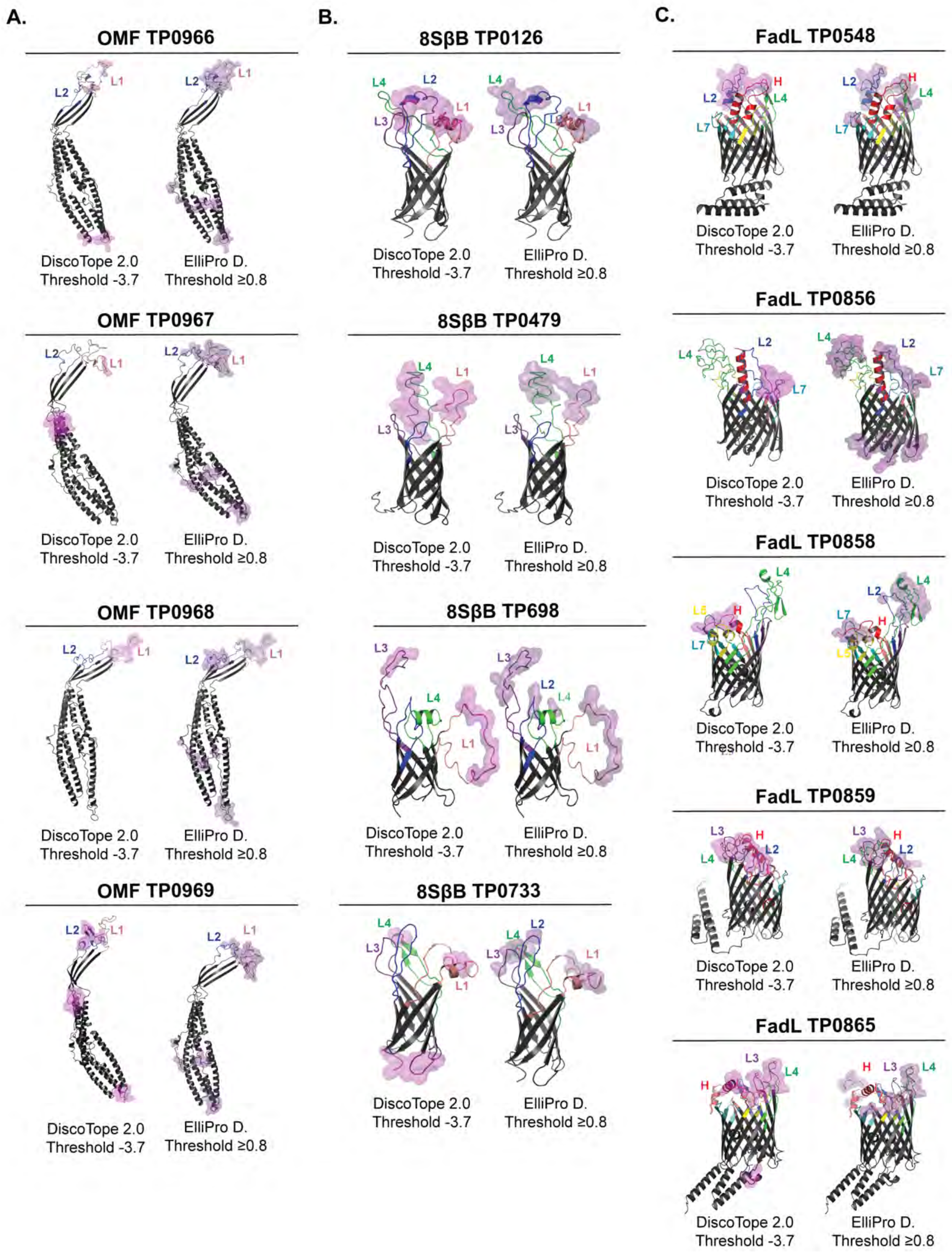
A.
OMF TP0966
DiscoTope 2.0
Threshold -3.7
ElliPro D.
Threshold ≥0.8
OMF TP0967
DiscoTope 2.0
Threshold -3.7
ElliPro D.
Threshold ≥0.8
OMF TP0968
DiscoTope 2.0
Threshold -3.7
ElliPro D.
Threshold ≥0.8
OMF TP0969
DiscoTope 2.0
Threshold -3.7
ElliPro D.
Threshold ≥0.8
B.
8SβB TP0126
DiscoTope 2.0
Threshold -3.7
ElliPro D.
Threshold ≥0.8
8SβB TP0479
DiscoTope 2.0
Threshold -3.7
ElliPro D.
Threshold ≥0.8
8SβB TP698
DiscoTope 2.0
Threshold -3.7
ElliPro D.
Threshold ≥0.8
8SβB TP0733
DiscoTope 2.0
Threshold -3.7
ElliPro D.
Threshold ≥0.8
C.
FadL TP0548
DiscoTope 2.0
Threshold -3.7
ElliPro D.
Threshold ≥0.8
FadL TP0856
DiscoTope 2.0
Threshold -3.7
ElliPro D.
Threshold ≥0.8
FadL TP0858
DiscoTope 2.0
Threshold -3.7
ElliPro D.
Threshold ≥0.8
FadL TP0859
DiscoTope 2.0
Threshold -3.7
ElliPro D.
Threshold ≥0.8
FadL TP0865
DiscoTope 2.0
Threshold -3.7
ElliPro D.
Threshold ≥0.8

**S4 Fig.**

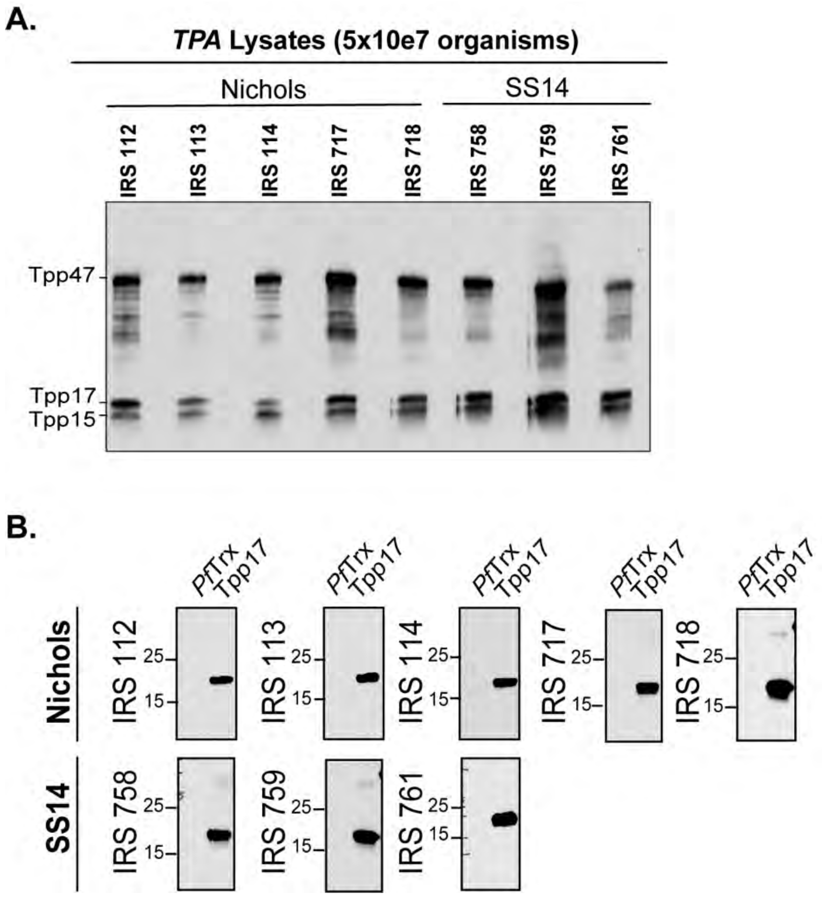
A.
TPA Lysates (5x10e7 organisms)
Nichols
SS14
IRS 112
IRS 113
IRS 114
IRS 717
IRS 718
IRS 758
IRS 759
IRS 761
Tpp47
Tpp17
Tpp15
B.
PfTrx Tpp17
Nichols
SS14
IRS 112
IRS 113
IRS 114
IRS 717
IRS 718
IRS 758
IRS 759
IRS 761
25
15

### S5 Fig.

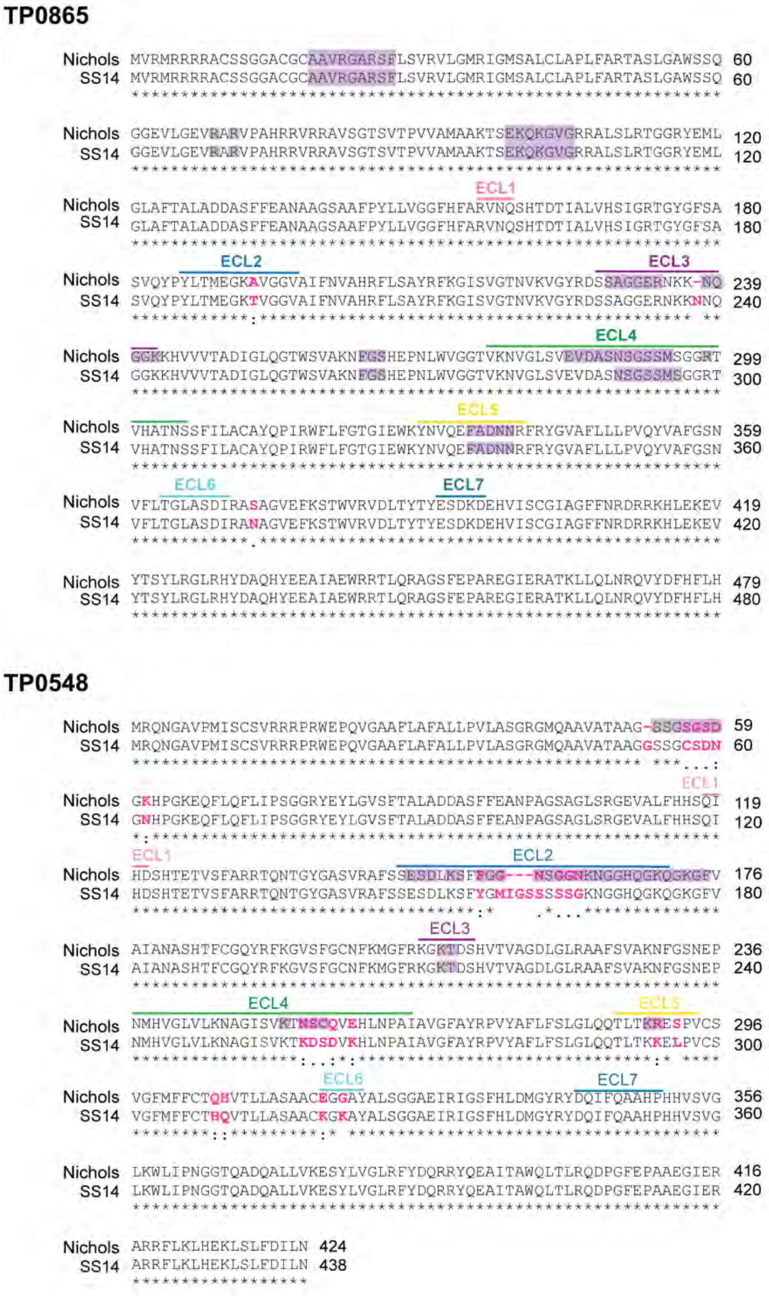
TP0865
Nichols
SS14
ECL1
ECL2
ECL3
ECL4
ECL5
ECL6
ECL7
60 120 180 239 299 359 419 479
60 120 180 240 300 360 420 480
TP0548
Nichols
SS14
ECL1
ECL2
ECL3
ECL4
ECL5
ECL6
ECL7
59 119 176 236 296 356 416 424
60 120 180 240 300 360 420 438

## TP0856

```
                                              ECL1
Nichols MVHYKSVFYKSAALVCGFVLAGASVAIASSEAAAKTRSKMSEFKRRAVSSPSGGRLSVLD 60
   SS14 MVHYKSVFYKSAALVCGFVLAGASVAIASSEAAAKTRSKMSEFKRRAVSSPSGGRLSVLD 60
        ************************************************************

Nichols GSFTALANDASFFEANPAGSANMTHSELTFAHTVGFNNSHAETLSYVGQSGNWGYGASMR 120
   SS14 GSFTALANDASFFEANPAGSANMTHSELTFAHTVGFNNSHAETLSYVGQSGNWGYGASMR 120
        ************************************************************
                  ECL2                                        ECL3
Nichols MFFPESGFNFSPSTGPVCTPASNPIKKLGGLGIVNFSRRFGGLSIGANLKAGFRDAQGLT 180
   SS14 MFFPESGFNFSPSTGPVCTPASNPIKKLGGLGIVNFSRRFGGLSIGANLKAGFRDAQGLT 180
        ************************************************************
                                                      ECL4
Nichols HLSLGTDVGLQWVGNVAKFFSSAEPNMYVGLSATNLGFTVKLPGSPFVLCRATGEQCCKT 240
   SS14 HLSLGTDVGLQWVGNVAKFFSSAEPNMYVGLSATNLGFTVKLPGSPFVLCRATGEQCCKT 240
        ************************************************************
                           ECL4
Nichols CSGRCTGVGTCCNGEKPCCKDCDCNCPCQDEATPGSPHATDTMLRAGFAYRPLSWFLFSV 300
   SS14 CSGRCTGVGTCCNGEKPCCKDCDCNCPCQDEATPGSPHATDTMLRAGFAYRPLSWFLFSV 300
        ************************************************************
              ECL5                                  ECL6
Nichols GVATRVNVSNLQVDHLNKRSSYALGMILDPVRFLTLLSGVAVNANGKVRAGVGAEIRVAC 360
   SS14 GVATRVNVSNLQVDHLNKRSSYALGMILDPVRFLTLLSGVAVNANGKVRAGVGAEIRVAC 360
        ************************************************************
                  ECL7
Nichols FQVSASYRYDSTGDEQQGTPHNMSLGASILLGRK 394
   SS14 FQVSASYRYDSTGDEQQGTPHNMSLGASILLGRK 394
        **********************************
```

## TP0858

```
Nichols MLRLPTARACITMGTMIRHTFTHRCGALLCALALGSSTMAATAAAKPKKGQMQKLRQRPV 60
   SS14 MLRLPTARACITMGTMIRHTFTHRCGALLCALALGSSTMAATAAAKPKKGQMQKLRQRPV 60
        ************************************************************
                                                      ECL1
Nichols WAPTGGRYASLDGAFTALANDASFFEANPAGSANMTHGELAFFHTTGFGSFHAETLSYVG 120
   SS14 WAPTGGRYASLDGAFTALANDASFFEANPAGSANMTHGELAFFHTTGFGSFHAETLSYVG 120
        ************************************************************
                               ECL2
Nichols QSGNWGYGASMRMFFPESGFDFSTTTEPVCTPASNPIKQRGAIGIINFARRIGGLSLGAN 180
   SS14 QSGNWGYGASMRMFFPESGFDFSTTTEPVCTPASNPIKQRGAIGIINFARRIGGLSLGAN 180
        ************************************************************
            ECL3                                            ECL4
Nichols LKAGFRDAQGLQHTSVSSDIGLQWVGNVAKSFTSEEPNLYIGLAATNLGLTVKVSDKIEN 240
   SS14 LKAGFRDAQGLQHTSVSSDIGLQWVGNVAKSFTSEEPNLYIGLAATNLGLTVKVSDKIEN 240
        ************************************************************
                           ECL4
Nichols CTSTCEKCGCCKERCCCNGKKACCKDCDCNCPCQDCNDKGTVHATDTMLRAGFAYRPFSW 300
   SS14 CTSTCEKCGCCKERCCCNGKKACCKDCDCNCPCQDCNDKGTVHATDTMLRAGFAYRPFSW 300
        ************************************************************
                          ECL5                                ECL6
Nichols FLFSLGATTSMNVQTLASSDAKSLYQNLAYSIGAMFDPFSFLSLSSSFRINHKANMRVGV 360
   SS14 FLFSLGATTSMNVQTLASSDAKSLYQNLAYSIGAMFDPFSFLSLSSSFRINHKANMRVGV 360
        ************************************************************
                            ECL7
Nichols GAEARIARIKLNAGYRCDVSDISSGSGCTGAKASHYLSLGGAILLGRN 408
   SS14 GAEARIARIKLNAGYRCDVNDISSGSGCTGAKASHYLSLGGAILLGRN 408
        *******************.****************************
```

## TP0859

```
Nichols LVRRPCVSAAPVRVGGRLVFGFARVGSRGLCLGALLLSPRIVLAQHVADAPLGARGVVPR 60
   SS14 MVRRPCVSAAPVRVGGRLVFGFARVGSRGLCLGALLLSPRIVLAQHVADAPLGARGVVPR 60
        :***********************************************************

Nichols SSLPRRTRAARATTLRSRGGVVSSRASGGTLVVTAQKPKVMARNDVDYRPLSLQAGGRQG 120
   SS14 SSLPRRTRAARATTLRSRGGVVSSRASGGTLVVTAQKPKVMARNDVDYRPLSLQAGGRQG 120
        ************************************************************
                                              ECL1
Nichols SLDLVATATADDASFFEANAAGSATIPRMTLAFFHTMRISDSHIDVLSFVGRAGRTGYGV 180
   SS14 SLDLVATATADDASFFEANAAGSATIPRMTLAFFHTMRISDSHIDVLSFVGRAGRTGYGV 180
        ************************************************************
            ECL2                                              ECL3
Nichols SARAFYPDMSSKTTGFVGIFNVSHAFSSAYRFKGVSVGANLKVGYRHTRGGGSSQSKSSN 240
   SS14 SARAFYPDMSSKTTGFVGIFNVSHAFSSAYRFKGVSVGANLKVGYRHTRGGGSSQSKSSN 240
        ************************************************************
                                                         ECL4
Nichols GKENHHIVLTADVGVRGAWTVSKNFGAHEPNLWAGVAFRNIGASINATNLHGNNGAGGSG 300
   SS14 GKENHHIVLTADVGVRGAWTVSKNFGAHEPNLWAGVAFRNIGASINATNLHGNNGAGGSG 300
        ************************************************************
            ECL4                                        ECL5
Nichols GGGGGNGDGKPAHVTDSRVILALAYQPVRYFLFGAGLEWLYNVGSIKAVNSLRYGAAFML 360
   SS14 GGGGGNGDGKPAHVTDSRVILALAYQPVRYFLFGAGLEWLYNVGSIKAVNSLRYGAAFML 360
        ************************************************************
                     ECL6                                  ECL7
Nichols FPLRQLAFSSSVVMKGMGPQQVRASAGAEVQFSHVRCTASYSYLWSATPTRPHYVSIGVA 420
   SS14 FPLRQLAFSSSVVMKGMGPQQVRASAGAEVQFSHVRCTASYSYLWSATPTRPHYVSIGVA 420
        ************************************************************

Nichols GFLKPVPEQPLWQEVYRSYLRGLRHYHAQRYAEAIAEWKRTLQQGVSFEPAREGIERATK 480
   SS14 GFLKPVPEQPLWQEVYRSYLRGLRHYHAQRYAEAIAEWKRTLQQGVSFEPAREGIERATK 480
        ************************************************************

Nichols LLQLNQKVHDFNIFT 495
   SS14 LLQLNQKVHDFNIF- 494
        **************
```

S6 Fig.

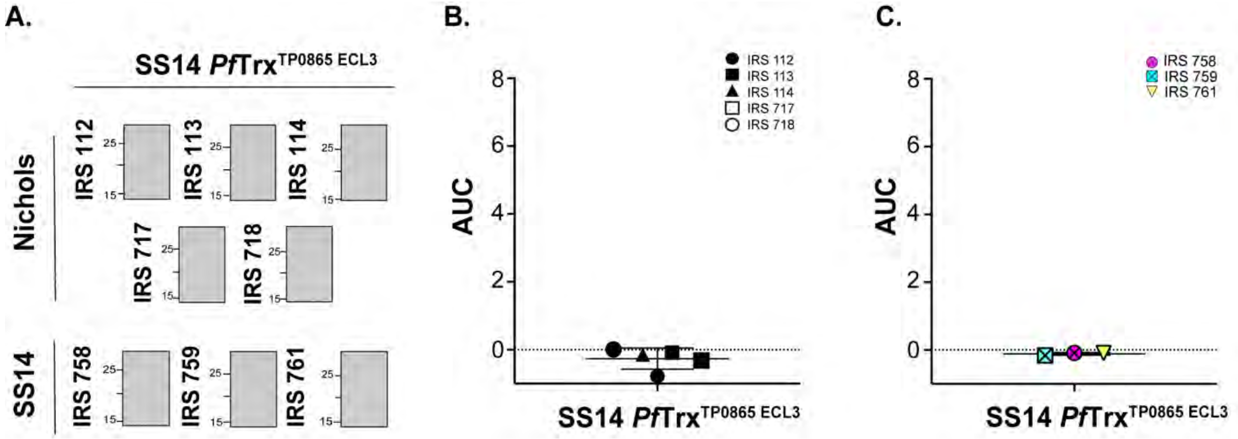
A.
SS14 PfTrx TP0865 ECL3
Nichols
SS14
IRS 112
IRS 113
IRS 114
IRS 717
IRS 718
IRS 758
IRS 759
IRS 761
25
15
B.
AUC
SS14 PfTrx TP0865 ECL3
IRS 112
IRS 113
IRS 114
IRS 717
IRS 718
8
6
4
2
0
C.
AUC
SS14 PfTrx TP0865 ECL3
IRS 758
IRS 759
IRS 761

**S7 Fig.**

**A.**

```
                       ********:** :* ***********: *.
TP0856 119 MRMFFPESGFNFSPSTGPVCTPASNPIKKLGGLGIVNF 156
TP0858 133 --MFFPESGFDFSTTTEPVCTPASNPIKQRGA------ 162
```

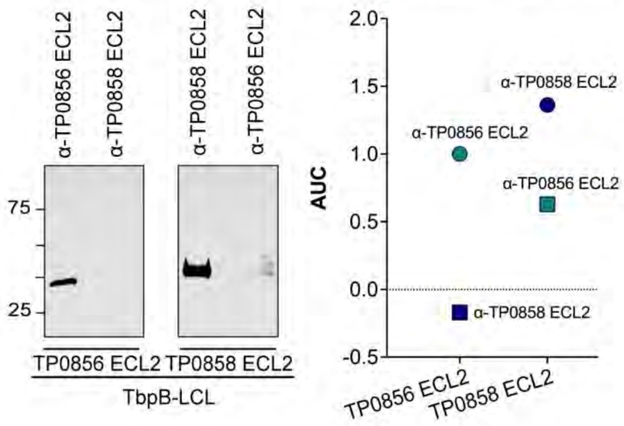


**B.**

```
               *  .     . *:* .**.   *     ****
TP0856 222 ---LPGSPFVLCRATGEQCCKTCS-GRCTGVGTCCNG 254
TP0858 224 AATNLGLTVK-VSDKIENCTSTCEKCGCCKERCCCNG 259
           :* *************.   *: ****
TP0856 255 EKPCCKDCDCNCPCQDEATPGSPHATD--- 281
TP0858 260 KKACCKDCDCNCPCQDCNDKGTVHATDTML 289
```

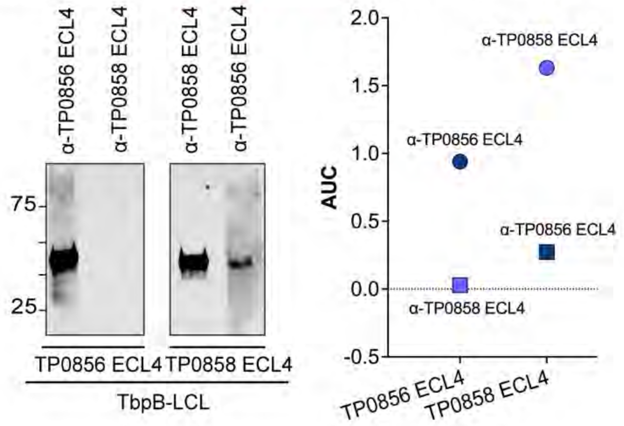
