## Supplemental Tables for "Immunodominant extracellular loops of *Treponema pallidum* FadL outer membrane proteins elicit antibodies with opsonic and growth-inhibitory activities"

### S1 Table

| Family | Protein | ECL | Name | Amino Acid Sequence | Primers |
|---|---|---|---|---|---|
| OMF | TP0966 | ECL1 | *Pf*Trx$^{TP0966/ECL1}$ | NLNGNGAGGMANGTPTLSPYVHLFFPTYQNLSLKADIAIKT | FW - GGGATGCGGTGGATCCCTGAATGGAACGGGGCC<br>RV - GACGACACGGGGATCCGGTCTTGATCGCAATATCCGC |
| | TP0966 | ECL2 | *Pf*Trx$^{TP0966/ECL2}$ | EGKSLKRILANVRMDFPGGGFWLGLNLPYPQWSRVE | FW - GGGATGCGGTGGATCCCTGAATGGAACGGGGCC<br>RV - GACGACACGGGGATCCGGTCTTGATCGCAATATCCGC |
| | TP0967 | ECL1 | *Pf*Trx$^{TP0967/ECL1}$ | AWNADGVKFRITPKASVAFPSFYNLTTHFGMTVTQPNGAAGGGGGGGGGDWQKT | FW - GGGATGCGGTGGATCCCTGAATGGAACGGGGCC<br>RV - GACGACACGGGGATCCGGTCTTGATCGCAATATCCGC |
| | TP0967 | ECL2 | *Pf*Trx$^{TP0967/ECL2}$ | TLDEGTGELSVAFPSVKITSALAIGYTGTLSI | FW - GGGATGCGGTGGATCCCTGAATGGAACGGGGCC<br>RV - GACGACACGGGGATCCGGTCTTGATCGCAATATCCGC |
| | TP0968 | ECL1 | *Pf*Trx$^{TP0968/ECL1}$ | IRGHSAGDFGLPRFGIKPIIGVRSPRYNNLVVSIDTARVTSIGNIS | FW - GGGATGCGGTGGATCCCTGAATGGAACGGGGCC<br>RV - GACGACACGGGGATCCGGTCTTGATCGCAATATCCGC |
| | TP0968 | ECL2 | *Pf*Trx$^{TP0968/ECL2}$ | MGTRKTFILFKGDGTESLEGSGTVALHMPSVNAQVEVKVPYAERGKHSRDKVG | FW - GGGATGCGGTGGATCCCTGAATGGAACGGGGCC<br>RV - GACGACACGGGGATCCGGTCTTGATCGCAATATCCGC |
| | TP0969 | ECL1 | *Pf*Trx$^{TP0969/ECL1}$ | LFNKKNGNGANGYKVEMAPHLSIASPFIGNSRLNLVAPRKLDGVTSTSTV | FW - GGGATGCGGTGGATCCCTGAATGGAACGGGGCC<br>RV - GACGACACGGGGATCCGGTCTTGATCGCAATATCCGC |
| | TP0969 | ECL2 | *Pf*Trx$^{TP0969/ECL2}$ | TDGEENKQLTNGMAPAAPSTTSTYGGTFNMAFPGGDSSFTVQNSKGLAGI | FW - GGGATGCGGTGGATCCCTGAATGGAACGGGGCC<br>RV - GACGACACGGGGATCCGGTCTTGATCGCAATATCCGC |
| 8SβB | TP0126 | ECL1 | *Pf*Trx$^{TP0126/ECL1}$ | PLFQVDWCNSGRGDDRNANAQTNGHKYIYPAFS | FW - GGGATGCGGTGGATCCCCGCTGTTCCAGGTTGATTGG<br>RV - GACGACACGGGGATCCGCTAAACGCCGGATAGATATACTTG |
| | TP0126 | ECL2 | *Pf*Trx$^{TP0126/ECL2}$ | SVQYHCSYPNNTYSPTTPYYYLAIPV | FW - GGGATGCGGTGGATCCAGCGTTCAGTATCACTGCAGC<br>RV - GACGACACGGGGATCCCACCGGAATCGCCAGATAATAG |
| | TP0126 | ECL3 | *Pf*Trx$^{TP0126/ECL3}$ | QHYYTSTYYGL | FW - CACCTATTATGGTCTGGGATCCCCGTGTCGTCTG<br>RV - CTGGTGTAATAGTGCTGACCGCATCCCGGAATGCT |
| | TP0126 | ECL4 | *Pf*Trx$^{TP0126/ECL4}$ | ATYSGVPRSCEKIIEEDRQQTNTRTAQF | FW - GAAGATCGTCAGCAAACCAATACCCGTACCGCGCAGTTTGGATCCCCGTGTCGTCTG<br>RV - CTCAATGATTTTTTCGCAGCTACGCGGAACGCCGCTGTAACCGCATCCCGGAATGCT |
| | TP0479 | ECL1 | *Pf*Trx$^{TP0479/ECL1}$ | VGAHPWGKEPAPRTDVLLYTP | FW - AGAACCGGCGCCGCGTACCGACGTTCTGCTGTATACCCCGTCCCCGTGTCGTCTGGTTG<br>RV - CGCGGCGCCGGTTCTTTACCCCACGGATGCGCACCAACTCCACCGCATCCCGGAATG |
| | TP0479 | ECL2 | *Pf*Trx$^{TP0479/ECL2}$ | AGLAFLMFRA | FW - GTTCCTGATGTTTCGTGGATCCCCGTGTCGTCTG<br>RV - GCCAGACCCGCATCCAGACCGCATCCCGGAATGCT |
| | TP0479 | ECL3 | *Pf*Trx$^{TP0479/ECL3}$ | LSTQDDKLVGIP | FW - CGATAAGCTGGTTGGTGGATCCCCGTGTCGTCTG<br>RV - TCTTGGGTGCTCAGAATACCGCATCCCGGAATGCT |
| | TP0479 | ECL4 | *Pf*Trx$^{TP0479/ECL4}$ | GSGVNVPLTKNLKKGANGNGQDLTWSNIWAKYCCTSLVM | FW - GGGATGCGGTGGATCCGGCAGCGGTGTGAACGTTCC<br>RV - GACGACACGGGGATCCCATAACCAGGCTGGTGCAGC |
| | TP0698 | ECL1 | *Pf*Trx$^{TP0698/ECL1}$ | GITSVYQFGSNGGSDGTSSGKGVSFDRLIGRVD | FW - GGGATGCGGTGGATCCGGTATTACCAGCGTGTACCAATTCG<br>RV - GACGACACGGGGATCCATCCACACGACCGATCAGACG |
| | TP0698 | ECL2 | *Pf*Trx$^{TP0698/ECL2}$ | SSLTNVFVRA | FW - TGACCAATGTCTTTGTGCGTGCATCCCCGTGTCGTCTGGTTG<br>RV - CAAAGACATTGGTCAACGAACTTCCACCGCATCCCGGAATG |
| | TP0698 | ECL3 | *Pf*Trx$^{TP0698/ECL3}$ | GVNICGDSCATSEGKSSAWYSKLLYSVPLN | FW - GGGATGCGGTGGATCCGGTGTTAACATTTGCGGTGACAGC<br>RV - GACGACACGGGGATCCGTTCAGCGGCACGCTATACAGC |
| | TP0698 | ECL4 | *Pf*Trx$^{TP0698/ECL4}$ | STAVGVRDFNFKEFTLPLSL | FW - ATTTCAACTTTAAGGAGTTTACCCTGCCGCTGAGCCTGTCCCCGTGTCGTCTGGTTG<br>RV - CCTTAAAGTTGAAATCACGAACGCCCACCGCGGTGCTTCCACCGCATCCCGGAATG |
| | TP0733 | ECL1 | *Pf*Trx$^{TP0733/ECL1}$ | GCQLYIAGGNGTNGSSSSGTNGNGNGKLLGGG | FW - GGGATGCGGTGGATCCTACATAGCAGGGGGAAATGGAACC<br>RV - GACGACACGGGGATCCCCCGCCCCCGAGCAGTTT |
| | TP0733 | ECL2 | *Pf*Trx$^{TP0733/ECL2}$ | FECYRTTGSNYYFSVPI | FW - GACCGGGTCAAACTATTACTTTTCTGTTCCCATCTCCCCGTGTCGTCTGGTTG<br>RV - TAGTTTGACCCGGTCGTGCGGTAACACTCAAATCCACCGCATCCCGGAATG |
| | TP0733 | ECL3 | *Pf*Trx$^{TP0733/ECL3}$ | LNIQSYLSKKAPGLI | FW - GAAGGCGCCGGGGCTTGGATCCCCGTGTCGTCTG<br>RV - TTGCTGAGATAGGACTGACCGCATCCCGGAATGCT |
| | TP0733 | ECL4 | *Pf*Trx$^{TP0733/ECL4}$ | YTQLGDIASSPDKCRAVGLA | FW - TGCAAGCTCCCCCGACAAGTGCAGAGCCGTGGGCCTTTCCCCGTGTCGTCTGGTTG<br>RV - TCGGGGGAGCTTGCAATGTCCCCAAGCTGCGTGTATCCACCGCATCCCGGAATG |
| FadL | TP0548 | ECL1 | *Pf*Trx$^{TP0548/ECL1}$ | QIHDSH | FW - AGATTCACGACAGCCACTCCCCGTGTCGTCTGGTTG<br>RV - GGCTGTCGTGAATCTGTCCACCGCATCCCGGAATG |
| | TP0548 | ECL2 | *Pf*Trx$^{TP0548/ECL2}$ | FSSESDLKSFFGGNSGGNKNGGHQGKQGKGFVAIA | FW - GGGATGCGGTGGATCCAGCGAGAGCGATCTGAAGAGC<br>RV - GACGACACGGGGATCCCGCAATCGCCACGAAGCC |
| | TP0548 | ECL3 | *Pf*Trx$^{TP0548/ECL3}$ | MGFRKGKTDSHVTV | FW - TAAGGGCAAAACCGACAGCCACGTGACCTCCCCGTGTCGTCTGGTTG<br>RV - TCGGTTTTGCCCTTACGGAAACCCATTCCACCGCATCCCGGAATG |
| | TP0548 | ECL4 | *Pf*Trx$^{TP0548/ECL4}$ | KNAGISVKTNSCQVEHLNPA | FW - AAACCAACAGCTGCCAGGTTGAGCACCTGAACCCGTCCCCGTGTCGTCTGGTTG<br>RV - GGCAGCTGTTGGTTTTCACGCTGATGCCCGCGTTTCCACCGCATCCCGGAATG |
| | TP0548 | ECL5 | *Pf*Trx$^{TP0548/ECL5}$ | QTLTKRESPVC | FW - GACCAAACGTGAAAGCCCGGTGTGCTCCCCGTGTCGTCTGGTTG<br>RV - CTTTCACGTTTGGTCAGGGTTTGTCCACCGCATCCCGGAATG |
| | TP0548 | ECL6 | *Pf*Trx$^{TP0548/ECL6}$ | ACEGGAYA | FW - GCGAGGGTGGCGCGTACGCGTCCCCGTGTCGTCTGGTTG<br>RV - ACGCGCCACCCTCGCACGCTCCACCGCATCCCGGAATG |
| | TP0548 | ECL7 | *Pf*Trx$^{TP0548/ECL7}$ | YDQIFQAAHPH | FW - GATCTTTCAGGCGGCGCACCCGCACTCCCCGTGTCGTCTGGTTG<br>RV - GCCGCCTGAAAGATCTGATCATATCCACCGCATCCCGGAATG |
| | TP0548 | Hatch | *Pf*Trx$^{TP0548/Hatch}$ | SGRGMQAAVATAAGSSGSGSDGKHPGKEQFLQFLIPSGGRYEYLGVSFTALADDASFFEANPAGSAGLSRG | FW - GGGATGCGGTGGATCCAGCGGCCGTGGTATGCAGG<br>RV - GACGACACGGGGATCCGCTCGGGATCAGAAATTGCAGG |
| | TP0856 | ECL1 | *Pf*Trx$^{TP0856/ECL1}$ | AHTVGFNNSHAETLSYV | FW - TAACAACAGCCACGCGGAGACCCTGAGCTACGTTGGATCCCCGTGTCGTCTGGT<br>RV - GCGTGGCTGTTGTTAAAGCCCACGGTGTGCGCGGATCCACCGCATCCCGG |
| | TP0856 | ECL2 | *Pf*Trx$^{TP0856/ECL2}$ | MRMFFPESGFNFSPSTGPVCTPASNPIKKLGGLGIVNF | FW - GGGATGCGGTGGATCCATGCGTATGTTCTTTCCGGAAAGC<br>RV - GACGACACGGGGATCCGAAGTTAACAATACCCAGGCCACCC |
| | TP0856 | ECL3 | *Pf*Trx$^{TP0856/ECL3}$ | GFRDAQGLTHLSLG | FW - CAGACCCTGCGCATCACGGAAACCGGATCCACCGCATCCCGG<br>RV - GATGCGCAGGGTCTGACCCACCTGAGCGGATCCCCGTGTCGTCTGGT |
| | TP0856 | ECL4 | *Pf*Trx$^{TP0856/ECL4}$ | LGFTVKLPGSPFVLCRATGEQCCKTCSGRCTGVGTCCNGEKPCCKDCDCNCPCQDEATPGSPHATDTML | FW - GGGATGCGGTGGATCCCTGCCGGGCAGCCCG<br>RV - GACGACACGGGGATCCCAGCATGGTATCGGTCGCGT |
| | TP0856 | ECL5 | *Pf*Trx$^{TP0856/ECL5}$ | TRVNVSNLQVDHLWKRS | FW - AGCGACGCGAAGAGCCTGTACCAAAACCTGGCGTATAGCGGATCCCCGTGTCGTCTGGTTGAACG<br>RV - GCTCTTCGCGTCGCTGCTCGCCAGGGTCTGAACGTTCATGCTGGTGGATCCACCGCATCCCGGAATGCTAAA |
| | TP0856 | ECL6 | *Pf*Trx$^{TP0856/ECL6}$ | AVNANGKVRAGV | FW - AGTGCGTGCGGGTGTTGGATCCCCGTGTCGTCTGGT<br>RV - ACACCCGCACGCACTTTGCCGTTCGCGTTAACGGATCCACCGCATCCCGG |
| | TP0856 | ECL7 | *Pf*Trx$^{TP0856/ECL7}$ | DSTGDEQQGTPHN | FW - ACAGCAAGGTACCCCGCACAACGGATCCCCGTGTCGTCTGGT<br>RV - GGGGTACCTTGCTGTTCATCACCGGTGCTGTCGGATCCACCGCATCCCGG |
| | TP0856 | Hatch | *Pf*Trx$^{TP0856/Hatch}$ | SSEAAAKTRSKMSEFKRRAVSSPSGGRLSVLDGSFTALANDASFFEANPAGSANMTH | FW - GCGGCGGCGAAGACCCGTAGCAAAATGGGATCCCCGTGTCGTCTGGT<br>RV - GGTCTTCGCCGCCGCCTCGCTGCTGGATCCACCGCATCCCGG |
| | TP0858 | ECL1 | *Pf*Trx$^{TP0858/ECL1}$ | FHTTGFGSFHAE | FW - GCTGCCGAAACCGGTGGTGTGAAAGGATCCACCGCATCCCGG<br>ACCGGTTTCGGCAGCTTTCACGCGGAAGGATCCCCGTGTCGTCTGGT |
| | TP0858 | ECL2 | *Pf*Trx$^{TP0858/ECL2}$ | MFFPESGFDFSTTTEPVCTPASNPIKQRGA | FW - GGGATGCGGTGGATCCAACTGGGGTTATGGCGCGAG<br>RV - GACGACACGGGGATCCCGCACCACGTTGCTTGATCG |
| | TP0858 | ECL3 | *Pf*Trx$^{TP0858/ECL3}$ | GFRDAQGLQHTSV | FW - CAGACCCTGCGCGTCACGAAAGCCGGATCCACCGCATCCCGG<br>RV - GACGCGCAGGGTCTGCAACACACCAGCGGATCCCCGTGTCGTCTGGT |
| | TP0858 | ECL4 | *Pf*Trx$^{TP0858/ECL4}$ | AATNLGLTVKVSDKIENCTSTCEKCGCCKERCCCNGKKACCKDCDCNCPCQDCNDKGTVHATDTML | FW - GGGATGCGGTGGATCCGCGGCGACCAACCTGGG<br>RV - GACGACACGGGGATCCCAGCATGGTATCGGTCGCGT |
| | TP0858 | ECL5 | *Pf*Trx$^{TP0858/ECL5}$ | SMNVQTLASSDAKSLYQNLA | FW - CGAGCAGCGACGCGAAGAGCCTGTACCAAAACTCCCCGTGTCGTCTGGTTG<br>RV - TCGCGTCGCTGCTCGCCAGGGTCTGAACGTTTCCACCGCATCCCGGAATG |

| | | | | | |
|---|---|---|---|---|---|
| | TP0858 | ECL6 | *Pf*Trx$^{\text{TP0858/ECL6}}$ | SFRINHKANMRV | FW - CGCCTTGTGGTTAATACGAAAGCTGGATCCACCGCATCCCGG<br>RV - ATTAACCACAAGGCGAACATGCGTGTGGGATCCCCGTGTCGTCTGGT |
| | TP0858 | ECL7 | *Pf*Trx$^{\text{TP0858/ECL7}}$ | RCDVSDISSGSGCTGAKASHY | FW - GCAGCCGCTACCGCTGCTGATATCGCTCACGTCGCAACGGGATCCACCGCATCCCGG<br>RV - AGCGGTAGCGGCTGCACCGGTGCGAAAGCGAGCCACTATGGATCCCCGTGTCGTCTGGT |
| | TP0858 | Hatch | *Pf*Trx$^{\text{TP0858/Hatch}}$ | AAAKPKKGQMQKLRQRPVWAPTGGRYASLDGAFTALANDASFFEANPAGSANMTH | FW - GGTCAAATGCAAAAACTGCGTCAACGTCCGGTGGGATCCCCGTGTCGTCTGGTTGAACG<br>RV - GTTTTTGCATTTGACCTTTCTTCGGCTTCGCCGCCGCGGATCCACCGCATCCCGGAATGCTAAA |
| | TP0859 | ECL1 | *Pf*Trx$^{\text{TP0859/ECL1}}$ | MRISDSH | FW - TGCGTATCAGCGACAGCTCCCCGTGTCGTCTGGTTG<br>RV - TGTCGCTGATACGCATTCCACCGCATCCCGGAATG |
| | TP0859 | ECL2 | *Pf*Trx$^{\text{TP0859/ECL2}}$ | FYPDMSSKTTGFV | FW - TGAGCAGCAAGACCACCGGCTTCGTGTCCCCGTGTCGTCTGGTTG<br>RV - TGGTCTTGCTGCTCATGTCCGGATATCCACCGCATCCCGGAATG |
| | TP0859 | ECL3 | *Pf*Trx$^{\text{TP0859/ECL3}}$ | GYRHTRGGGSSQSKSSNGKENHHIVLT | FW - GGGATGCGGTGGATCCTATCGTCACACCCGTGGT<br>RV - GACGACACGGGGATCCCACAATGTGGTGGTTCTCTTTACCG |
| | TP0859 | ECL4 | *Pf*Trx$^{\text{TP0859/ECL4}}$ | FRNIGASINATNLHGNNGAGGSGGGGGGNGDGKPAHVTDS | FW - GGGATGCGGTGGATCCTTTCGTAACATCGGTGCGAGC<br>RV - GACGACACGGGGATCCGCTATCGGTCACGTGCGC |
| | TP0859 | ECL5 | *Pf*Trx$^{\text{TP0859/ECL5}}$ | LYNVGSIKAVNSL | FW - GGGCAGCATTAAGGCGGTTAACAGCCTGTCCCCGTGTCGTCTGGTTG<br>RV - GCCTTAATGCTGCCCACGTTGTACAGTCCACCGCATCCCGGAATG |
| | TP0859 | ECL6 | *Pf*Trx$^{\text{TP0859/ECL6}}$ | VMKGMGPQQVR | FW - AGGCATGGGTCCGCAACAAGTTCGTTCCCCGTGTCGTCTGGTTG<br>RV - TGCGGACCCATGCCTTTCATAACTCCACCGCATCCCGGAATG |
| | TP0859 | ECL7 | *Pf*Trx$^{\text{TP0859/ECL7}}$ | YLWSATPTRPHY | FW - GCGCGACCCCGACCCGTCCGCACTACTCCCCGTGTCGTCTGGTTG<br>RV - GGGTCGGGGTCGCGCTCCACAGATATCCACCGCATCCCGGAATG |
| | TP0859 | Hatch | *Pf*Trx$^{\text{TP0859/Hatch}}$ | QHVADAPLGARGVVPRSSLPRRTRAARATTLRSRGGVVSSRASGGTLVVTAQKPKVMARNDVDYRPLSLQAG GRQGSLDLVATATADDASFFEANAAGSATIPR | FW - GGGATGCGGTGGATCCCAACATGTTGCGGATGCGC<br>RV - GACGACACGGGGATCCCAGGCTCAGCGGACGATAATC |
| | TP0865 | ECL1 | *Pf*Trx$^{\text{TP0865/ECL1}}$ | ARVNQSH | FW - CCCGGGTTAATCAATCGTCCCCGTGTCGTCTGGTTG<br>RV - ATTGATTAACCCGGGCTCCACCGCATCCCGGAATG |
| | TP0865 | ECL2 | *Pf*Trx$^{\text{TP0865/ECL2}}$ | QYPYLTMEGKAVGGVAI | FW - AATGGAGGGAAAGGCAGTAGGCGGCGTTGCCATTTCCCCGTGTCGTCTGGTTG<br>RV - GCCTTTCCCTCCATTGTCAGGTAGGGATACTGTCCACCGCATCCCGGAATG |
| | TP0865 | ECL3 | *Pf*Trx$^{\text{TP0865/ECL3}}$ | GYRDSSAGGERNKKNQGGKKHVVV | FW - GGAGAGGAATAAAAAAAACCAGGGGGGTAAGAAGTCCCCGTGTCGTCTGGTTG<br>RV - TTTTTATTCCTCTCCCCGCCGGCTGAAGAATCTCCACCGCATCCCGGAATG |
| | TP0865 | ECL4 | *Pf*Trx$^{\text{TP0865/ECL4}}$ | VKNVGLSVEVDASNSGSSMSGGRTVHATNSS | FW - GGGATGCGGTGGATCCGTCAAAAACGTTGGACTTTCAGTTG<br>RV - GACGACACGGGGATCCACTCGAGTTTGTAGCGTGCAC |
| | TP0865 | ECL5 | *Pf*Trx$^{\text{TP0865/ECL5}}$ | KYNVQEFADNNRF | FW - GCAAGAATTTGCAGACAACAATAGATTCTCCCCGTGTCGTCTGGTTG<br>RV - TCTGCAAATTCTTGCACGTTGTACTTTCCACCGCATCCCGGAATG |
| | TP0865 | ECL6 | *Pf*Trx$^{\text{TP0865/ECL6}}$ | LTGLASDI | FW - CACCGGGCTCGCCTCAGATTCCCCGTGTCGTCTGGTTG<br>RV - GAGGCGAGCCCGGTGAGTCCACCGCATCCCGGAATG |
| | TP0865 | ECL7 | *Pf*Trx$^{\text{TP0865/ECL7}}$ | YESDKDE | FW - TGAGTCAGACAAGGATGAGTCCCCGTGTCGTCTGGTTG<br>RV - TCCTTGTCTGACTCATATCCACCGCATCCCGGAATG |
| | TP0865 | Hatch | *Pf*Trx$^{\text{TP0865/Hatch}}$ | RTASLGAWSSQGGEVLGEVRARVPAHRRVRRAVSGTSVTPVVAMAAKTSEKQKGVGRRALSLRTGGRYEMLG LAFTALADDASFFEANAAGSAAFPY | FW - GGGATGCGGTGGATCCTCTTCGCAAGGGGGTGAAGTTCTGG<br>RV - GACGACACGGGGATCCCCTCAGCGAGAGGGCACG |
| ***Pf*Trx** | | | *Pf*Trx$^{\text{Empty}}$ | MGSSHHHHHHSSGLVPRGSHMSSGIIEYDGEIDFTKGRVVLWFSIPGCGGSPCRLVERFMTELSEYFEDIQIVHINA GKWKNIVDKFNILNVPTLVYLKDGREVGRQNLIRSKEEILKKLKELQE | FW - TGCAGGAGTAACTCGAGCACCACCACCA<br>RV - CGAGTTACTCCTGCAGCTCTTTCAGT |
| **FadL ECLs used for animal immunizations** | | | *Pf*Trx$^{\text{TP0856/ECL2}}$ No AviTag | MRMFFPESGFNFSPSTGPVCTPASNPIKKLGGLGIVNF | FW - TGCAGGAGTAACTCGAGCACCACCACCA<br>RV - CGAGTTACTCCTGCAGCTCTTTCAGT |
| | | | *Pf*Trx$^{\text{TP0856/ECL4}}$ No AviTag | LPGSPFVLCRATGEQCCKTCSGRCTGVGTCCNGEKPCCKDCDCNCPCQDEATPGSPHATDTML | |
| | | | *Pf*Trx$^{\text{TP0858/ECL2}}$ No AviTag | MFFPESGFDFSTTTEPVCTPASNPIKQRGA | |
| | | | *Pf*Trx$^{\text{TP0858/ECL4}}$ No AviTag | AATNLGLTVKVSDKIENCTSTCEKCGCCKERCCCNGKKACCKDCDCNCPCQDCNDKGTVHATDTML | |
| | | | *Pf*Trx$^{\text{TP0865/ECL3}}$ No AviTag | DSSAGGERNKKNQGGKK | |
| **TbpB loopless C-lobe (TbpB-LCL)** | | | TbpB-LCL$^{\text{TP0856/ECL2}}$ | GSSSENKLTTVLDAVELTLNDKKIKNLDNFSNAAQLVVDGIMIDLAGTEFTRKFEHTMRMFFPESGFNFSPSTGPVC TPASNPIKKLGGLGIVNFTYEVEVCCSNLNYLKYGMLTRKGKQVEQSMFLQGERTDEKEIPTDQNVVYRGSWYGHI ANGTSWSGNASDKEGGNRAEFTVNFADKKITGKLTAENRQAQTFTIEGMIQGNGFEGTAKTAESGFDLDQKNTTRT PKAYITDAKVKGGFYGPKAEELGGWFAYPGNAPEGKQEKATVVFGAKRQQPVQGEAAAKEAAAKGLNDIFEAQKIE WHE | synthetic gene |
| | | | TbpB-LCL$^{\text{TP0856/ECL4}}$ | GSSSENKLTTVLDAVELTLNDKKIKNLDNFSNAAQLVVDGIMIDLAGTEFTRKFEHTLPGSPFVLCRATGEQCCKTC SGRCTGVGTCCNGEKPCCKDCDCNCPCQDEATPGSPHATDTMLTYEVEVCCSNLNYLKYGMLTRKGKQVEQSM FLQGERTDEKEIPTDQNVVYRGSWYGHIANGTSWSGNASDKEGGNRAEFTVNFADKKITGKLTAENRQAQTFTIEG MIQGNGFEGTAKTAESGFDLDQKNTTRTPKAYITDAKVKGGFYGPKAEELGGWFAYPGNAPEGKQEKATVVFGAK RQQPVQGEAAAKEAAAKGLNDIFEAQKIEWHE | synthetic gene |
| | | | TbpB-LCL$^{\text{TP0858/ECL2}}$ | GSSSENKLTTVLDAVELTLNDKKIKNLDNFSNAAQLVVDGIMIDLAGTEFTRKFEHTMFFPESGFDFSTTTEPVCTPA SNPIKQRGATYEVEVCCSNLNYLKYGMLTRKGKQVEQSMFLQGERTDEKEIPTDQNVVYRGSWYGHIANGTSWS GNASDKEGGNRAEFTVNFADKKITGKLTAENRQAQTFTIEGMIQGNGFEGTAKTAESGFDLDQKNTTRTPKAYITDA KVKGGFYGPKAEELGGWFAYPGNAPEGKQEKATVVFGAKRQQPVQGEAAAKEAAAKGLNDIFEAQKIEWHE | synthetic gene |
| | | | TbpB-LCL$^{\text{TP0858/ECL4}}$ | GSSSENKLTTVLDAVELTLNDKKIKNLDNFSNAAQLVVDGIMIDLAGTEFTRKFEHTAATNLGLTVKVSDKIENCTST CEKCGCCKERCCCNGKKACCKDCDCNCPCQDCNDKGTVHATDTMLTYEVEVCCSNLNYLKYGMLTRKGKQVE QSMFLQGERTDEKEIPTDQNVVYRGSWYGHIANGTSWSGNASDKEGGNRAEFTVNFADKKITGKLTAENRQAQTF TIEGMIQGNGFEGTAKTAESGFDLDQKNTTRTPKAYITDAKVKGGFYGPKAEELGGWFAYPGNAPEGKQEKATVVF GAKRQQPVQGEAAAKEAAAKGLNDIFEAQKIEWHE | synthetic gene |
| | | | TbpB-LCL$^{\text{TP0865/ECL3}}$ | GSSSENKLTTVLDAVELTLNDKKIKNLDNFSNAAQLVVDGIMIDLAGTEFTRKFEHTDSSAGGERNKKNQGGKKTYE VEVCCSNLNYLKYGMLTRKGKQVEQSMFLQGERTDEKEIPTDQNVVYRGSWYGHIANGTSWSGNASDKEGGNR AEFTVNFADKKITGKLTAENRQAQTFTIEGMIQGNGFEGTAKTAESGFDLDQKNTTRTPKAYITDAKVKGGFYGPKAE ELGGWFAYPGNAPEGKQEKATVVFGAKRQQPVQGEAAAKEAAAKGLNDIFEAQKIEWHE | synthetic gene |
| | | | TbpB-LCL$^{\text{Empty}}$ | GSSSENKLTTVLDAVELTLNDKKIKNLDNFSNAAQLVVDGIMIDLAGTEFTRKFEHTINGKTYEVEVCCSNLNYLKYG MLTRKGKQVEQSMFLQGERTDEKEIPTDQNVVYRGSWYGHIANGTSWSGNASDKEGGNRAEFTVNFADKKITGKL TAENRQAQTFTIEGMIQGNGFEGTAKTAESGFDLDQKNTTRTPKAYITDAKVKGGFYGPKAEELGGWFAYPGNAPE GKQEKATVVFGAKRQQPVQGEAAAKEAAAKGLNDIFEAQKIEWHE | synthetic gene |
| | | | TbpB-LCL$^{\text{ECL}}$ into pRB1B vector | | FW - CGCGCGGCAGCCATATGGGCAGCAGCAGCGAAAAC<br>RV - GGTGGTGGTGCTCGAGTTATTCATGCCATTCAATTTTCTGC |

Amino acid sequences of ECL boundaries predicted by trRosetta (highlighted in bold) and AlphFold3 (underlined) for all three OMP families investigated in this study. Amino acid sequences and primer pairs utilized for *Pf*Trx-scaffolded ECL constructs (ECL1-Salmon, ECL2-Blue, ECL3-Purple, ECL4-Green, ECL5-Yellow, ECL6-Cyan, ECL7-Dark Teal, and Hatch-Red).

### S2 Table

| | Attached *TPA* Rabbit Assay | | | | | | | | | | | |
|---|---|---|---|---|---|---|---|---|---|---|---|---|
| | 1% | | | | 5% | | | | 10% | | | |
| | **Mean % (Range %)** | ***p*-value$^{\#}$** | ***p*-value$^{*}$** | ***p*-value$^{\blacktriangle}$** | **Mean % (Range %)** | ***p*-value$^{\#}$** | ***p*-value$^{*}$** | ***p*-value$^{\blacktriangle}$** | **Mean % (Range %)** | ***p*-value$^{\#}$** | ***p*-value$^{*}$** | ***p*-value$^{\blacktriangle}$** |
| **TpCM-2 medium** | 83 (81-84) | n.s | 0.0002 | < 0.0001 | 75 (71-78) | n.s | n.s | n.s | 83 (81-84) | n.s | n.s | n.s |
| **NRS** | 82 (81-82) | n.s | 0.0003 | < 0.0001 | 81 (78-83) | n.s | 0.0191 | 0.0278 | 82 (81-82) | n.s | n.s | n.s |
| **IRS 112** | 31 (26-43) | < 0.0001 | 0.0189 | n.s | 39 (30-44) | 0.0002 | n.s | n.s | 26 (13-39) | < 0.0001 | n.s | 0.0244 |
| **α-TP0856 ECL2** | 52 (45-59) | 0.0008 | | n.s | 52 (50-55) | 0.0245 | | n.s | 52 (38-66) | n.s | | n.s |
| **α-TP0856 ECL4** | 88 (87-89) | n.s | < 0.0001 | < 0.0001 | 86 (83-91) | n.s | 0.0026 | 0.0037 | 75 (69-78) | n.s | n.s | n.s |
| **α-TP0858 ECL2** | 79 (65-89) | n.s | 0.0011 | < 0.0001 | 68 (50-79) | n.s | n.s | n.s | 58 (49-69) | n.s | n.s | n.s |
| **α-TP0858 ECL4** | 74 (64-79) | n.s | 0.0189 | 0.0002 | 66 (51-89) | n.s | n.s | n.s | 67 (51-80) | n.s | n.s | n.s |
| **α-TP0865 ECL3** | 94 (92-96) | n.s | < 0.0001 | < 0.0001 | 90 (89-91) | n.s | 0.0007 | 0.0010 | 74 (66-85) | n.s | n.s | n.s |
| **α-BamA ECL4** | 43 (35-49) | < 0.0001 | n.s | | 53 (49-60) | 0.0356 | n.s | | 60 (38-76) | n.s | n.s | |
| **α-TP0751** | 84 (83-85) | n.s | < 0.0001 | < 0.0001 | 95 (92-97) | n.s | 0.0001 | 0.0002 | 70 (69-72) | n.s | n.s | n.s |
| **α-Tpp17** | 80 (78-82) | | 0.0008 | < 0.0001 | 80 (78-82) | | 0.0245 | 0.0356 | 83 (82-84) | | n.s | n.s |
| **α-*Pf*Trx** | 87 (86-88) | n.s | < 0.0001 | < 0.0001 | 93 (91-95) | n.s | 0 | 0.0003 | 91 (89-91) | n.s | 0.0071 | n.s |
| | Statistical analysis was done using one-way ANOVA. $^{\#}$: vs. α-Tpp17; $^{*}$: vs. α-TP0856 ECL2; $^{\blacktriangle}$: vs. α-BamA ECL4 | | | | | | | | | | | |

### S3 Table

| | **Attached *TPA* Mouse Assay** | | | | | | | |
|---|---|---|---|---|---|---|---|---|
| | **5%** | | | | | | | |
| | **Mean % (Range %)** | ***p*-value#** | ***p*-value*** | **p-value△** | **p-value♠** | ***p*-value★** | ***p*-value✦** | ***p*-value▲** |
| **TpCM-2 medium** | 84 (83-85) | n.s | 0.0008 | 0.0131 | < 0.0001 | < 0.0001 | < 0.0001 | < 0.0001 |
| **NMS** | 84 (82-86) | n.s | 0.0008 | 0.0131 | < 0.0001 | < 0.0001 | < 0.0001 | < 0.0001 |
| **MSS** | 52 (49-56) | < 0.0001 | n.s | 0.0055 | 0.0036 | 0.0015 | n.s | n.s |
| **α-TP0856 ECL2** | 64 (54-72) | < 0.0001 | | n.s | < 0.0001 | < 0.0001 | 0.0019 | 0.0006 |
| **α-TP0856 ECL4** | 68 (61-76) | < 0.0001 | n.s | | < 0.0001 | < 0.0001 | 0.0001 | < 0.0001 |
| **α-TP0858 ECL2** | 34 (34-35) | < 0.0001 | < 0.0001 | < 0.0001 | | n.s | n.s | n.s |
| **α-TP0858 ECL4** | 33 (31-34) | < 0.0001 | < 0.0001 | < 0.0001 | n.s | | n.s | n.s |
| **α-TP0865 ECL3** | 46 (42-48) | < 0.0001 | 0.0019 | 0.0001 | n.s | n.s | | n.s |
| **α-BamA ECL4** | 45 (42-45) | < 0.0001 | 0.0006 | < 0.0001 | n.s | n.s | n.s | |
| **α-TP0751** | 90 (84-93) | n.s | < 0.0001 | 0.0003 | < 0.0001 | < 0.0001 | < 0.0001 | < 0.0001 |
| **α-Tpp17** | 93 (91-94) | | < 0.0001 | < 0.0001 | < 0.0001 | < 0.0001 | < 0.0001 | < 0.0001 |
| **α-*Pf*Trx** | 86 (80-92) | n.s | 0.0002 | 0.0029 | < 0.0001 | < 0.0001 | < 0.0001 | < 0.0001 |

Statistical analysis was done using one-way ANOVA. #: vs. α-Tpp17; *: vs. α-TP0856 ECL2; △: vs. α-TP0856 ECL4; ♠: vs. α-TP0858 ECL2; ★: vs. α-TP0858 ECL4; ✦: vs. α-TP0865 ECL3; ▲: vs. α-BamA ECL4

### S4 Table

**Statistical Significance among *in vivo* OMPs**

| | p value# |
|---|---|
| **OMP** | TP0858 |
| TP0966 | <0.0001 |
| TP0967 | n.s |
| TP0968 | <0.0001 |
| TP0969 | <0.0001 |
| TP0126 | <0.0001 |
| TP0479 | <0.0001 |
| TP0698 | <0.0001 |
| TP0733 | 0.0002 |
| TP0548 | <0.0001 |
| TP0856 | 0.0009 |
| TP0858 | |
| TP0859 | <0.0001 |
| TP0865 | <0.0001 |
| TP0326 | <0.0001 |
| TP0751 | n.s |
| TP0435 | n.s |

**Statistical Significance among *in vitro* OMPs**

| | p value# | | | | |
|---|---|---|---|---|---|
| **OMP** | TP0967 | TP0733 | TP0858 | TP0751 | TP0435 |
| TP0966 | n.s | n.s | <0.0001 | <0.0001 | <0.0001 |
| TP0967 | | n.s | 0.0008 | n.s | <0.0001 |
| TP0968 | 0.0003 | <0.0001 | <0.0001 | <0.0001 | <0.0001 |
| TP0969 | <0.0001 | <0.0001 | <0.0001 | <0.0001 | <0.0001 |
| TP0126 | n.s | 0.0002 | <0.0001 | <0.0001 | <0.0001 |
| TP0479 | <0.0001 | <0.0001 | <0.0001 | <0.0001 | <0.0001 |
| TP0698 | <0.0001 | <0.0001 | <0.0001 | <0.0001 | <0.0001 |
| TP0733 | n.s | | n.s | n.s | <0.0001 |
| TP0548 | 0.0006 | <0.0001 | <0.0001 | <0.0001 | <0.0001 |
| TP0856 | 0.0002 | <0.0001 | <0.0001 | <0.0001 | <0.0001 |
| TP0858 | 0.0008 | n.s | | n.s | 0.0007 |
| TP0859 | n.s | n.s | <0.0001 | <0.0001 | <0.0001 |
| TP0865 | <0.0001 | <0.0001 | <0.0001 | <0.0001 | <0.0001 |
| TP0326 | <0.0001 | <0.0001 | <0.0001 | <0.0001 | <0.0001 |
| TP0751 | n.s | n.s | n.s | | <0.0001 |
| TP0435 | <0.0001 | <0.0001 | 0.0007 | <0.0001 | |

Statistical analysis was done using two-way ANOVA #: *p* value of ≦ 0.001
